# Presynaptic autophagy is coupled to the synaptic vesicle cycle via ATG-9

**DOI:** 10.1101/2020.12.28.424508

**Authors:** Sisi Yang, Daehun Park, Laura Manning, Sarah E. Hill, Mian Cao, Zhao Xuan, Ian Gonzalez, Lin Shao, Ifechukwu Okeke, Pietro De Camilli, Daniel A. Colón-Ramos

## Abstract

Autophagy is a cellular degradation pathway essential for neuronal health and function. Autophagosome biogenesis occurs at synapses, is locally regulated and increases in response to neuronal activity. The mechanisms that couple autophagosome biogenesis to synaptic activity remain unknown. In this study we determine that trafficking of ATG-9, the only transmembrane protein in the core autophagy pathway, links the synaptic vesicle cycle with autophagy. ATG-9 positive vesicles in *C. elegans* are generated from the trans-Golgi network via AP3-dependent budding, and delivered to presynaptic sites. At presynaptic sites, ATG-9 undergoes exo-endocytosis in an activity-dependent manner. Mutations that disrupt endocytosis, including one associated with Parkinson’s disease, result in abnormal ATG-9 accumulation at clathrin-rich synaptic foci and defects in activity-dependent presynaptic autophagy. Our findings uncover regulated key steps of ATG-9 trafficking at presynaptic sites, and provide evidence that ATG-9 exo-endocytosis couples autophagosome biogenesis at presynaptic sites with the activity-dependent synaptic vesicle cycle.

**Highlights:**

- In *C. elegans*, ATG-9 is delivered to presynaptic sites in vesicles generated from the trans-Golgi network via AP-3-dependent budding
- ATG-9 vesicles undergo activity-dependent exo-endocytosis at presynaptic sites
- Mutations in endocytic proteins, including a mutation associated with Parkinson’s disease, result in abnormal ATG-9 accumulation at clathrin-rich foci
- Abnormal accumulation of ATG-9 at clathrin-rich foci is associated with defects in activity-dependent presynaptic autophagy

## Introduction

Macroautophagy (herein called autophagy) is an evolutionarily conserved cellular degradative process that is essential for neuronal physiology and survival (Son et al., 2012, Stavoe and Holzbaur, 2019, Azarnia Tehran et al., 2018, Kulkarni et al., 2018, Liang and Sigrist, 2018, Menzies et al., 2017, Vijayan and Verstreken, 2017, Menzies et al., 2015, Tsukada and Ohsumi, 1993, Yorimitsu and Klionsky, 2005). Neurons are particularly vulnerable to dysfunctional organelles and damaged proteins due to their post-mitotic nature, their polarized morphology and their high metabolic activity states during neuronal stimulation. Autophagy is regulated to cater to these neurophysiological needs. For example, local autophagosome biogenesis occurs near synapses and autophagosome biogenesis is coupled to the neuronal activity state (Bunge, 1973, Soukup et al., 2016, Maday et al., 2012, Stavoe et al., 2016, Katsumata et al., 2010, Shehata et al., 2012, Hill et al., 2019). Disruption of synaptic autophagy has been associated with the accumulation of damaged proteins and organelles, synaptic dysfunction and neurodegenerative diseases, including Parkinson’s disease (Hoffmann et al., 2019, Hill et al., 2019, Lynch-Day et al., 2012, Zavodszky et al., 2014, Karabiyik et al., 2017, Cheung and Ip, 2009).

Molecules that regulate synaptic transmission and function, including proteins involved in synaptic vesicle exo-endocytosis, were reported to regulate autophagy at presynaptic sites (Soukup et al., 2016, George et al., 2016, Vanhauwaert et al., 2017, Murdoch et al., 2016, Kononenko et al., 2017, Binotti et al., 2015). For example, in *Drosophila*, endophilin A, a protein mainly known for its role in endocytosis, was proposed to directly regulate autophagosome formation by inducing curved membranes that can recruit autophagic machinery (Soukup et al., 2016, Milosevic et al., 2011). Synaptojanin 1, a phosphoinositide phosphatase implicated in the endocytic recycling of synaptic vesicles (Cremona et al., 1999, Verstreken et al., 2003, Harris et al., 2000), was also reported to play roles in the control of synaptic autophagy in zebrafish and *Drosophila* (Vanhauwaert et al., 2017, George et al., 2016). Recent studies have revealed links between these canonical endocytic proteins and early-onset parkinsonism (EOP), suggesting a relationship between the synaptic vesicle cycle (which is tied to synaptic activity), autophagy and neurodegenerative diseases (Vidyadhara et al., 2019, Trinh and Farrer, 2013, Alegre-Abarrategui and Wade-Martins, 2009, Bandres-Ciga et al., 2019, Schreij et al., 2016, Quadri et al., 2013, Krebs et al., 2013). Yet, the mechanistic links underlying the coupling between synaptic activity and autophagosome formation remain unknown.

In this study we examined the dynamics of ATG-9, the only transmembrane protein of the core autophagy machinery, at synapses of *C. elegans* and mammalian neurons. ATG-9 is thought to promote local autophagosome biogenesis through its role as a lipid scramblase that cooperates with the lipid transport protein ATG2 in the nucleation of the isolation membrane in nascent autophagosomes (Karanasios et al., 2016, Reggiori et al., 2005, Reggiori et al., 2004, Sawa-Makarska et al., 2020, Guardia et al., 2020, Matoba et al., 2020, Matoba and Noda, 2020, Maeda et al., 2020, Gomez-Sanchez et al., 2018). We find that at synapses, ATG-9 links the synaptic vesicle cycle to autophagy. Specifically, we observe that in *C. elegans* neurons, ATG-9 is delivered to presynaptic sites in vesicles generated by the trans-Golgi network (TGN) via AP-3-dependent budding. At presynaptic sites, ATG-9 positive vesicles undergo exo-endocytosis in a synaptic activity-dependent manner. Mutants that disrupt synaptic endocytic traffic, including a *synaptojanin 1/unc-26* allele that mimics a Parkinson disease mutation, result in abnormal accumulation of ATG-9 in clathrin-rich foci, and defects in activity-dependent synaptic autophagy. Mutations that affect autophagosome biogenesis also result in abnormal accumulations of ATG-9 in clathrin-rich foci, further suggesting a relation between endocytic trafficking of ATG-9 and nucleation of autophagosomes at presynaptic sites. In mammalian hippocampal neurons, mutations in endocytic proteins similarly result in abnormal ATG9 accumulation in nerve terminals, indicating conserved mechanisms of ATG-9 trafficking at synapses. Collectively our studies identify the regulated dynamics of ATG-9 trafficking at presynaptic sites and provide insight into mechanisms that couple the synaptic vesicle cycle (related to synaptic activity) to presynaptic autophagy.

## Results

### In *C. elegans* ATG-9 is transported to synapses in vesicles generated in the trans-Golgi network (TGN) via AP-3-dependent budding

Autophagy occurs at presynaptic sites in response to synaptic activity, and transmembrane protein ATG-9 plays a critical role in local synaptic autophagy (Stavoe et al., 2016, Hill et al., 2019, Soukup et al., 2016, Wang et al., 2015, Shehata et al., 2012). To understand the dynamics of ATG-9 at presynaptic sites, we first examined the *in vivo* localization of ATG-9 in the AIY interneurons of *C. elegans*. AIYs are a pair of bilaterally symmetric interneurons which display a stereotypical distribution of presynaptic specializations along their neurites ((White et al., 1986, Colon-Ramos et al., 2007); (Figures 1A-1B)). Simultaneous visualization of the presynaptic marker mCherry::RAB-3 and ATG-9::GFP revealed that ATG-9 localization in neurons is discrete, compartmentalized and enriched at subcellular structures in the cell body and at presynaptic regions ((Stavoe et al., 2016); (Figures 1C-1F, 1C’)).

**Fig 1.**
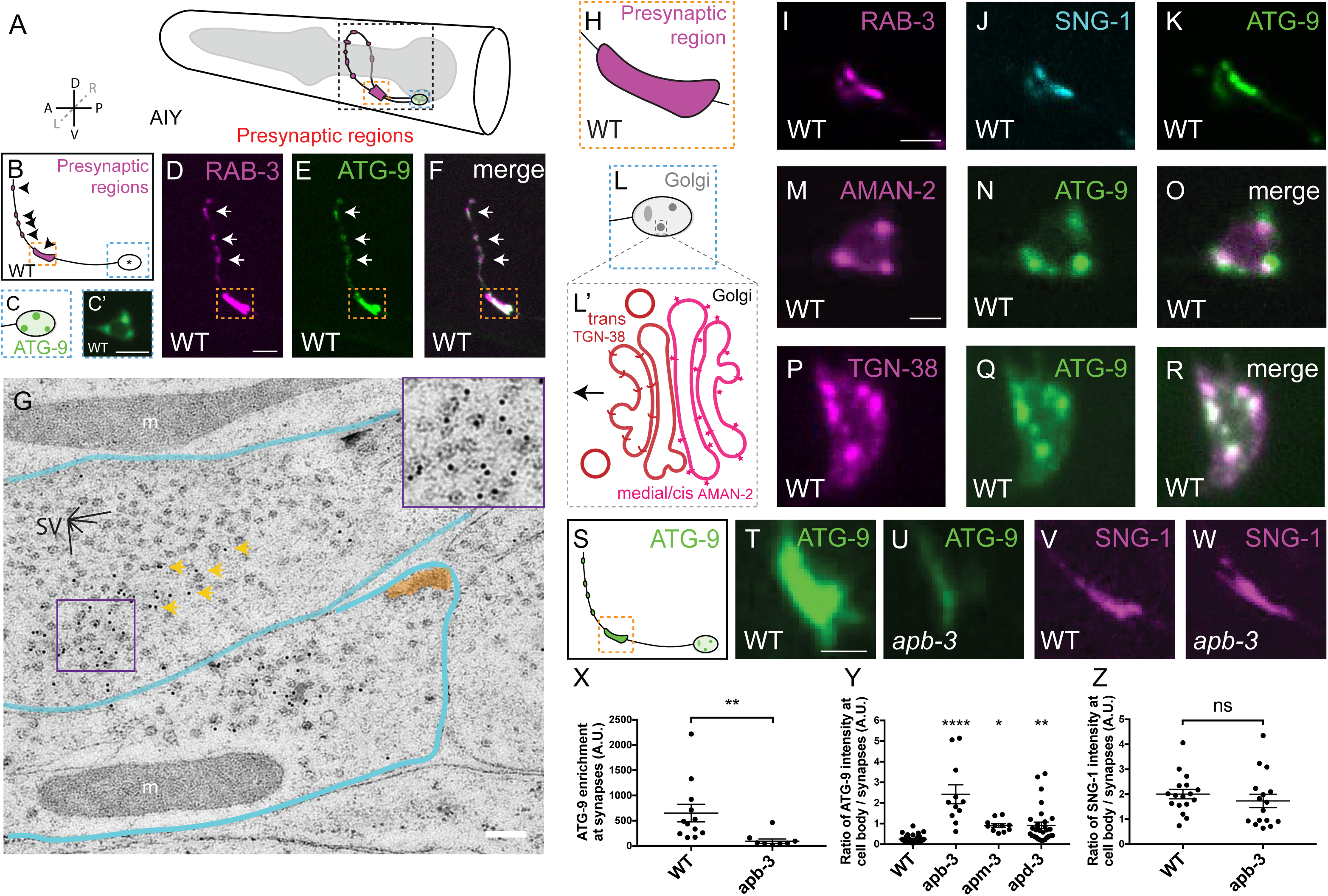
In *C. elegans* ATG-9 is transported to synapses in vesicles generated in the trans-Golgi network (TGN) via AP-3-dependent budding. (A) Schematic of the head of *C. elegans*, including pharynx (grey region) and the two bilaterally symmetric AIY interneurons (in black dashed box) with presynaptic regions (magenta). The synaptic-rich region is highlighted with an orange dashed square and cell bodies with a blue dashed square. In axis, A, anterior; P, posterior; L, left; R, right; D, dorsal; V, ventral. (B) Schematic of a single AIY interneuron with cell body (in blue dashed square) and presynaptic regions (magenta and black arrowheads, and synaptic-rich region highlighted with an orange dashed square). (C) Schematic of a cell body in the AIY interneurons, with ATG-9 localization represented in green. (C’) Representative confocal micrograph of ATG-9::GFP localization at the cell body of AIY in a wild-type animal (as in C, and blue dashed box in B). (D-F) Representative confocal micrographs of RAB-3::mCherry (pseudo-colored magenta) (D), ATG-9::GFP (E) and a merged channels (F) in the synaptic regions of a representative wild-type AIY interneuron. The arrows and the dashed box highlight the presynaptic specializations. (G) Immunogold electron microscopy of transgenic animals with panneuronal expression of ATG-9::GFP, with antibodies directed against GFP. Note that the majority of immunogold particles localize to the presynaptic areas occupied by synaptic vesicles, but not to all synaptic vesicle (see Supplementary Fig 1). Blue line, outline of neurons. Dense projections, shaded in orange. “m”, mitochondria. SV, examples of synaptic vesicles. Yellow arrows point to examples of immunogold particles. (H) Schematic of synaptic-rich region in the AIY interneurons (I-K) Representative confocal micrographs of RAB-3::mCherry (pseudo-colored magenta) (I), SNG-1::BFP (pseudo-colored cyan) (J) and ATG-9::GFP (K) at synaptic-rich region (corresponding to H, also orange dashed box in B) in wild type. (L) Schematic of the AIY interneuron cell body with Golgi labeled (as grey puncta), (L’) Schematic of the Golgi apparatus with medial/cis-Golgi-specific protein AMAN-2 (magenta) and trans-Golgi-specific protein TGN-38 (red). (M-O) Confocal micrographs of AMAN-2::GFP (pseudo-colored magenta) (M), ATG-9::mCherry (pseudo-colored green) (N) and merged channels (O) in the cell body of AIY. (P-R) Confocal micrographs of TGN-38::mCherry (pseudo-colored magenta) (P), ATG-9::GFP (Q) and merged channels (R) in the cell body of AIY. (S) Schematic of an AIY interneuron with ATG-9 (green). Pre-synaptic-rich region (Zone 2) is highlighted by orange rectangle. (T-U) Confocal micrographs of ATG-9::GFP at Zone 2 in wild type (T) and *apb-3(ok429)* mutants (U). (V-W) Confocal micrographs of SNG-1::GFP (pseudo-colored magenta) at Zone 2 in wild type (V) and *apb-3(ok429)* mutants (W). (X) Quantification of ATG-9::GFP enrichment at Zone 2 of AIY neurons in wild-type and *apb-3(ok429)* mutant animals. Error bars correspond to standard error of the mean (SEM). **p<0.01 by Welch’s t test between wild-type and mutant animals. Each dot in the scatter plot represents a single animal. (Y) Quantification of the ratio of ATG-9 intensity at cell body / synapses of AIY neurons in wild-type, *apb-3(ok429)*, *apm-3(gk771233)* and *apd-3(gk805642)* mutant animals. Error bars represent standard error of the mean (SEM). *p<0.05, **p<0.01 and ****p<0.0001 (between wild type and the mutants) by ordinary one-way ANOVA with Dunnett’s multiple comparisons test between wild-type and the mutant groups. Each dot in the scatter plot represents a single animal. (Z) Quantification of ratio of SNG-1 intensity at cell body / synapses of AIY neurons in wild-type and *apb-3(ok429)* mutant animals. “ns”: not significant (between wild type and the mutants) by Welch’s t test between wild-type and mutant animals. Each dot in the scatter plot represents a single animal. Scale bars 5μm in (C’); 5μm in (D) for (D)-(F); 200nm in (G); 2μm in (I) for (I)-(K); 2μm in (M) for (M)-(R); 2μm in (T) for (T)-(W).

To identify ATG-9 positive structures at synapses, we performed post-embedding immunogold electron microscopy of transgenic animals expressing ATG-9::GFP, by using antibodies directed against GFP. We observed that the majority of the immunogold particles (75%) localized to the presynaptic areas occupied by synaptic vesicles, with occasional localization of gold particles on the plasma membrane (<10%) (Figures 1G, S1A-S1D). However, the distribution of immunogold particles was generally non-homogeneous in the synaptic vesicle-positive areas. Accordingly, comparison by light microscopy with the localization of mCherry::RAB-3 and of a well-established synaptic vesicle integral membrane protein, SNG-1::BFP, revealed strong colocalization between mCherry::RAB-3 and SNG-1::BFP, but subtle differences between the colocalization of these two proteins and ATG-9::GFP (Figures 1H-1K), consistent with ATG-9 being enriched on a subpopulation of vesicles.

To determine the site of ATG-9 localization within the cell soma, we co-expressed either ATG-9::mCherry or ATG-9::GFP with other organelle markers (Reggiori et al., 2005, Karanasios et al., 2016, van der Vaart and Reggiori, 2010, Puri et al., 2013). We observed that ATG-9::GFP was concentrated at sites that overlapped with trans-Golgi marker TGN-38::mCherry and that ATG-9::mCherry was directly adjacent to the medial/cis-Golgi marker AMAN-2::GFP, but did not overlap with other organelles markers (Figures 1L-1R, 1L’, S1E-S1K). These findings, which are consistent with observations from yeast and mammalian culture cells (Noda, 2017, Webber et al., 2007, Ohashi and Munro, 2010), indicate a trans-Golgi-specific enrichment of ATG-9 in the neuronal cell body.

The trans-Golgi network (TGN) is where vesicles destined for transport to distinct subcellular locations are packaged. Coat protein complexes such as members of the heterotetrameric family of adaptor proteins (APs) regulate this process by sorting cargoes into distinct vesicular carriers (Park and Guo, 2014, Nakatsu and Ohno, 2003, Mattera et al., 2017, Badolato and Parolini, 2007, Dell’Angelica et al., 1997). In vertebrate, AP-4 is particularly important in exporting transmembrane protein ATG9A from the Golgi apparatus, leading to neurological disorders (Yamamoto et al., 2012, De Pace et al., 2018, Mattera et al., 2017). In *C. elegans* and other invertebrates, however, no orthologues of AP-4 have been characterized. Which proteins are then required for the biogenesis of ATG-9 positive vesicles?

We tested mutants for AP complexes UNC-101/AP-1, DPY-23/AP-2 and AP-3 (Park and Guo, 2014, Nakatsu and Ohno, 2003) and did not observe abnormal localization of ATG-9 in loss-of-function alleles of *unc-101(m1)/ap-1* or *dpy-23(e840)/ap-2* (data not shown). However, a putative null allele for *apb-3(ok429)*, a gene encoding a subunit of the AP-3 complex, displayed a reduction of ATG-9 at synapses (Figures 1S-1U, 1X). The AP-3 complex comprises four subunits - apb-3/β3-adaptin, apd-3/δ-adaptin, apm-3/μ3 and aps-3/σ3. Putative null alleles for the three AP-3 complex subunits (apb-3/β3-adaptin, apm-3/μ3, apd-3/δ-adaptin) resulted in enrichment of ATG-9 at the cell body (Figure 1Y). Interestingly, these alleles did not affect the localization of the synaptic vesicle proteins SNG-1 or RAB-3 to presynaptic sites, suggesting that the observed decrease of ATG-9 at synapses is not due to a general problem in synaptic vesicle biogenesis (Figures 1V-1W, 1Z, S1L-S1O). Together, our findings reveal that *in vivo* in *C. elegans* neurons, enrichment of ATG-9 at presynaptic sites results from AP-3 mediated export of ATG-9 positive vesicles at the TGN.

### ATG-9 undergoes exo-endocytosis at presynaptic sites

The concentration of ATG-9 on vesicles at presynaptic sites and its occasional localization in the axonal plasma membrane raised the possibility that this protein may be a component of vesicles that undergo exo-endocytosis. To address this possibility, we imaged ATG-9 synaptic localization in *C. elegans* neurons in mutants with disrupted exo-endocytic traffic at presynaptic sites (Harris et al., 2000, Watanabe et al., 2014, Sudhof, 1995, Saheki and De Camilli, 2012) (Figure 2A). We observed that endocytic mutants *unc-26(e205)/synaptojanin 1, unc-57(ok310)/endophilin A*, *unc-11(e47)/AP180* and temperature-sensitive *dyn-1(ky51)* displayed defects in ATG-9 localization at synapses (Figures 2B-2H). For example, in the AIY interneuron presynaptic-rich region (termed Zone 2; (Colon-Ramos et al., 2007)), the ATG-9::GFP signal is predominantly concentrated in a homogenous manner across this presynaptic area (Figures 1E, 1K, 2B-2B’, 2C). However, in all the endocytic mutants examined, most animals display abnormal accumulation of ATG-9::GFP into multiple subsynaptic foci (Figures 2D-2H, 2SA-B). To quantify the genetic expressivity of the phenotype, we defined an index for ATG-9 mislocalization (briefly, the index was defined as the number of local signal peaks divided by their width, see STAR Methods). We found that in the endocytic mutants there was a significant difference in the subsynaptic localization of ATG-9, with ATG-9 abnormally enriched at discrete foci within the presynaptic regions (Figure 2K).

**Fig 2.**
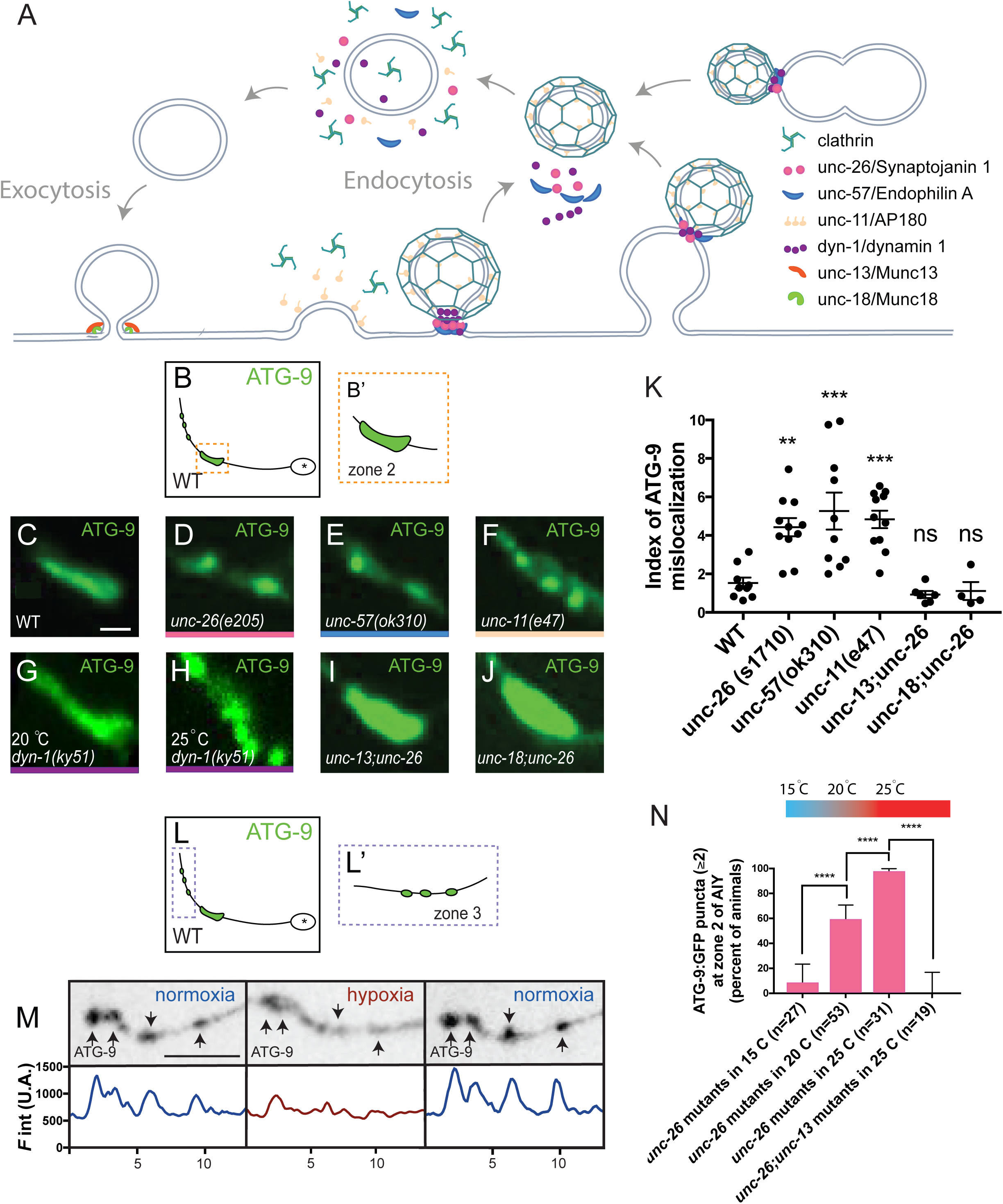
ATG-9 undergoes exo-endocytosis at presynaptic sites. (A) Schematic of the proteins required for the synaptic vesicle cycle examined in this study, with vertebrate and *C. elegans* gene names (Saheki and De Camilli, 2012, Watanabe et al., 2014, Gan and Watanabe, 2018). (B-B’’) Schematic of ATG-9 localization in AIY neurons (B), enlargement of the synaptic-rich region of Zone 2 (B’). (C-J) Confocal micrographs of ATG-9::GFP at AIY Zone 2 in wild type (C), *unc-26(e205)* (D), *unc-57(ok310)* (E) and *unc-11(e47)* (F) mutants, temperature-sensitive *dyn-1(ky51)* mutants in the permissive temperature 20°C (G), in the restrictive temperature 25°C (H), *unc-13(s69);unc-26(e205)* (I) and *unc-18(e81);unc-26(s1710)* (J) double mutants. (K) Quantification of the index of ATG-9 mislocalization (see STAR Methods) in wild type, *unc-26(s1710)*, *unc-57(ok310)*, *unc-11(e47)* mutants, *unc-13(s69);unc-26(e205)* and *unc-18(e81);unc-26(s1710)* double mutants. Error bars show standard error of the mean (SEM). “ns” (not significant), **p<0.01 and ***p<0.001 (between wild type and the mutants) by ordinary one-way ANOVA with Dunnett’s multiple comparisons test between wild-type and the mutant groups. Each dot in the scatter plot represents a single animal. (L, L’) Schematic of ATG-9 localization in AIY neurons (L), enlargement of the distal part of the neurite with synaptic clusters of Zone 3 (L’). (M) Confocal micrographs of ATG-9::GFP (pseudo-colored black) at AIY Zone 3 in *pfk-1.1*(*gk*9226*89*) mutants prehypoxia/normoxia (left panels), after 10min of transient hypoxia (middle panels) and another iteration of 10min of normoxia (right panels). Corresponding fluorescence intensity is shown below the line scan image. (N) Quantification of the percentage of animals displaying two (or more) ATG-9 subsynaptic foci at AIY Zone 2 in *unc-26(s*1710) mutants raised at indicated temperatures. Higher temperatures in AIY result in increased activity state, and autophagy (Clark et al., 2006, Hawk et al., 2018, Hill et al., 2019). *unc-13(s69);unc-26(e205)* double mutants were also raised at 25°C and quantified. The number of animals examined in each condition is indicated by “n”. Error bars show standard error of the mean (SEM). ****p<0.0001 by two-tailed Fisher’s exact test. Scale bars 2μm in (C) for (C)-(J); 5μm in (M)

Are endocytic proteins acting cell autonomously in neurons to regulate ATG-9 subsynaptic localization? To address this question, we focused on UNC-26/synaptojanin-1. Expressing *unc-26* cDNA cell specifically in the AIY interneurons of *unc-26(e205)* mutants rescued ATG-9 defects at the synapse, indicating that UNC-26 acts cell autonomously in neurons to prevent abnormal ATG-9 accumulation at subsynaptic foci (Figures S2C-S2E).

If the accumulation of ATG-9 at abnormal foci in endocytic mutants is due to abnormal endocytic traffic, then mutants in exocytosis should suppress this phenotype. To test this hypothesis, we examined putative null alleles *unc-13(s69)/Munc1*3 and *unc-18(e81)/Munc18*, which encode essential components of synaptic vesicle exocytosis (Hata et al., 1993, Richmond et al., 1999). Single mutants of *unc-13(s69)* and *unc-18(e81)* did not disrupt ATG-9 localization (Figures S2A, S2F-S2G). Importantly, double mutants of *unc-13(s69);unc-26(e205)* and *unc-18(e81);unc-26(s1710)* suppressed the *unc-26/synaptojanin 1* phenotype, indicating that the accumulation of ATG-9 at abnormal foci depends on synaptic vesicle exocytosis (Figures 2I-2K, S2A).

Our genetic perturbations in *C. elegans* are consistent with ATG-9 positive vesicles undergoing exo-endocytosis at presynaptic sites. To better examine this, we imaged ATG-9 dynamics in a loss-of-function allele of phosphofructokinase 1/*pfk-1.1*(*gk922689*). The absence of phosphofructokinase, like the absence of other glycolytic proteins, results in impaired synaptic vesicle endocytosis during transient hypoxia (Jang et al., 2016). Through the use of a microfluidic device that allows precise control of transient cycles of normoxia and hypoxia, we can temporally control the endocytic reaction (Jang et al., 2020). We examined ATG-9 localization in the synaptic Zone 3 region (Figure L-L’), in which we can observe discrete and interspersed presynaptic specializations (compare to the Zone 2 region (Figure 2B-B’), which consists of one large and continuous presynaptic area). Visualization of ATG-9 in the Zone 3 region enabled us to determine local ATG-9 dynamics due to transient inhibition of endocytosis. We observed that in *pfk-1.1*(*gk922689*) mutants, transient inhibition of endocytosis during transient hypoxia correlated with changes in ATG-9 localization: namely, ATG-9 relocalized from discrete presynaptic clusters in the synaptic Zone 3 region, to a more diffuse distribution, consistent with what would be expected if ATG-9 were trapped at the plasma membrane due to short-term defects in endocytosis. Conversely, removing the endocytic block by shifting to normoxia rescued the localization of ATG-9 to the presynaptic clusters (Figures 2L-2M). Together, our data indicate that ATG-9 positive vesicles undergo exo-endocytosis at presynaptic sites by using the synaptic vesicle cycling machinery.

### ATG-9 mislocalization phenotypes are enhanced under conditions that increase AIY activity-state

We next examined if the mislocalization phenotype of ATG-9 could be modified based on physiologically relevant stimuli known to increase the activity state of the neuron, and known to increase synaptic autophagy (Hill et al., 2019). AIY neurons in *C. elegans* are part of the thermotaxis circuit, which allows animals to navigate towards their cultivation temperature, and the activity state of AIY increases based on the cultivation temperature at which the organism is reared (Clark et al., 2006, Hawk et al., 2018). At higher cultivation temperatures, AIY displays increases in activity-dependent synaptic autophagy (Hill et al., 2019).

We observed that the penetrance of the ATG-9 phenotype in *unc-26(s1710)/synaptojanin 1* mutants similarly varied depending on the cultivation temperature of the animals. At higher cultivation temperatures, known to increase the activity state of AIY and synaptic autophagy, we observed a higher percentage of *unc-26(s1710)/synaptojanin 1* mutant animals displaying abnormal ATG-9 foci at synapses (Figure 2N). Moreover, temperature-dependent increases of abnormal ATG-9 foci in *unc-26(s1710)/synaptojanin* 1 mutant animals were suppressed by the exocytosis mutant *unc-13(s69)* (Figure 2N).

### Biochemical evidence that ATG9 travels to and from the plasma membrane

Our results are consistent with published evidence that ATG-9 can be exposed at the cell surface and then re-internalized by endocytosis in mammalian fibroblastic cells, as revealed by immunocytochemistry following perturbation of a critical endocytic factor, dynamin, either pharmacologically or by dominant interference (Puri et al., 2013, Feng and Klionsky, 2017, Zhou et al., 2017). To obtain direct biochemical evidence for ATG9 exo-endocytosis, we performed cell surface biotinylation experiments (with Sulfo-NHS-LC-Biotin) in tamoxifen-inducible dynamin 1 and 2 double knock-out (DKO) mouse fibroblasts (Ferguson et al., 2009). A pool of ATG9A, and of transferrin receptor as a control, was detected at the plasma membrane in control fibroblasts, and these pools were enhanced (Figures 3A-3C) in fibroblasts where the expression of dynamin 1 and 2 had been suppressed by tamoxifen (Figures S3A-S3B), providing biochemical evidence that ATG-9 travels to and from the plasma membrane, and that its internalization depends on endocytic proteins.

**Fig 3.**
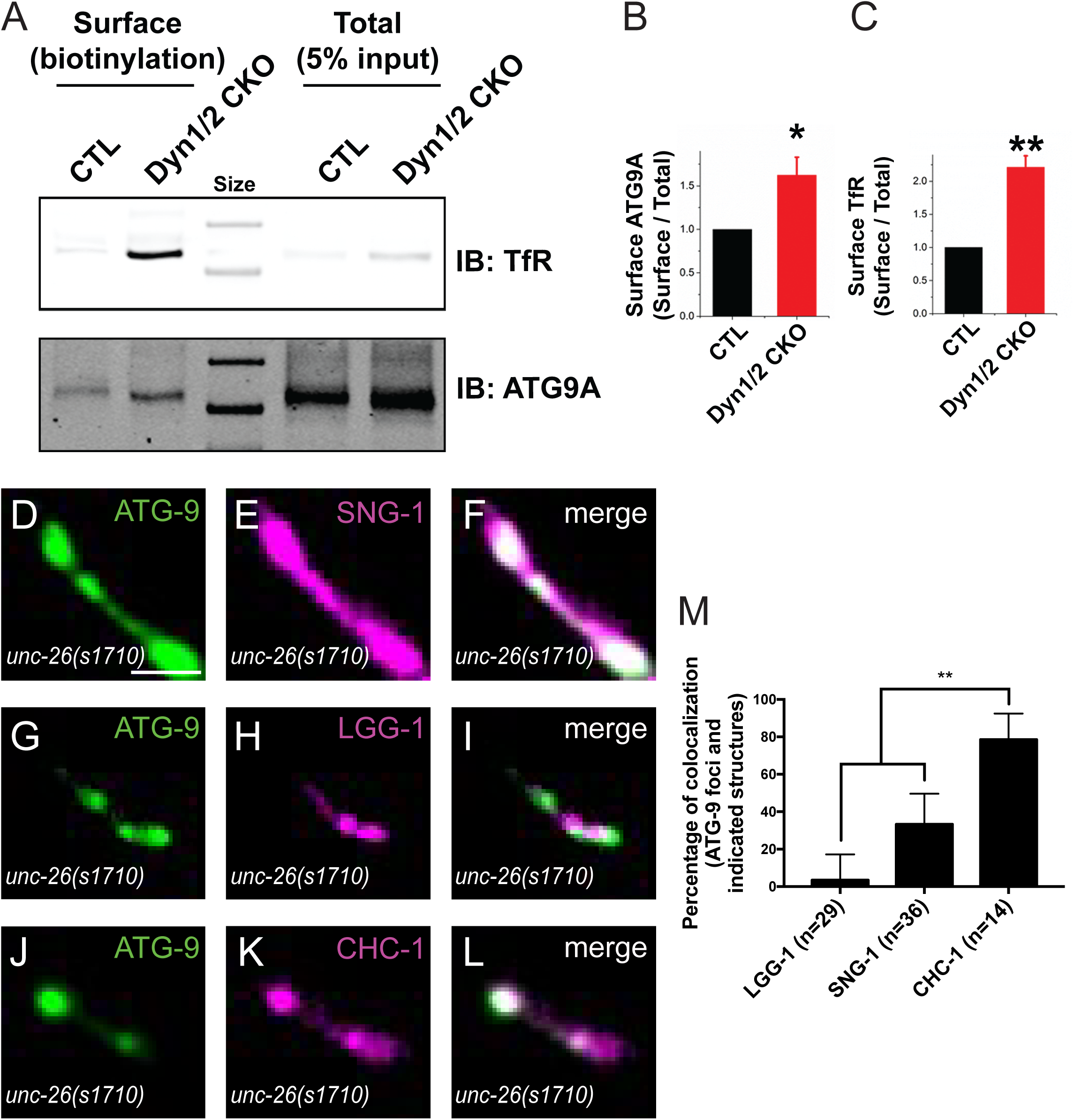
In *unc-26/synaptojanin 1* mutants, ATG-9 accumulates at presynaptic, clathrin-rich sites. (A-C) Surface levels of ATG9A and transferrin receptor (TfR) in control and dynamin 1 and 2 conditional double knock-out (Dyn1/2 CKO) fibroblasts. (A) Immunoblots (IB) for transferrin receptor (TfR) and ATG9A of total cell extracts and of material recovered by streptavidin affinity purification following surface biotinylation. (B and C) Quantification of surface / total levels of ATG9 and TfR in Dyn1/2 CKO fibroblasts relative to the control cells (CTL). Error bars show standard error of the mean (SEM). *p<0.05 and **p<0.01 by Student’s t test. (D-F) Confocal micrographs of ATG-9::GFP (D), SNG-1::BFP (pseudo-colored magenta) (E) and merged channels (F) at AIY Zone 2 in *unc-26(s*1710) mutants. (G-I) Confocal micrographs of ATG-9::mCherry (pseudo-colored green) (G), GFP::LGG-1 (pseudo-colored magenta) (H) and merged channels (I) in *unc-26(s1710)* mutants. (J-L) Confocal micrographs of ATG-9::GFP (J), BFP::clathrin heavy chain (CHC-1) (pseudo-colored magenta) (K) and merged (L) in *unc-26(s*1710) mutants. (M) Percentage of ATG-9 foci that colocalize with LGG-1, the synaptic-vesicle associated transmembrane protein Synaptogyrin/SNG-1 and CHC-1 puncta. The number of foci examined in each condition are indicated by the “n”. Error bars show standard error of the mean (SEM). **p<0.01 by two-tailed Fisher’s exact test. Scale bars 2μm in (D) for (D)-(L).

### In *unc-26/synaptojanin 1* mutants, ATG-9 accumulates at presynaptic, clathrin-rich sites

What are the subsynaptic foci where ATG-9 is enriched in endocytic mutants? We first examined if ATG-9 accumulated with synaptic vesicles proteins in such foci. Defects in the endocytic pathway result in the redistribution of synaptic vesicle membrane proteins to the plasma membrane and endocytic intermediates, which is reflected in a less clustered localization of intrinsic and peripheral synaptic vesicle proteins in neurites (Harris et al., 2000, Verstreken et al., 2003, Ferguson et al., 2007, Raimondi et al., 2011, Milosevic et al., 2011). Consistent with these findings, RAB-3::mCherry and SNG-1::GFP displayed diffuse localization in *unc-26(s1710)/synaptojanin 1* mutants (Figures S3C-S3F). These phenotypes are distinct from the ones observed for ATG-9, as SNG-1::BFP in *unc-26(s1710)* mutants failed to localize similarly to ATG-9::GFP at subsynaptic foci (Figures 3D-3F, 3M). Our data indicate that while ATG-9 undergoes exo-endocytosis at the synapse in an activity dependent manner, mutations in endocytic proteins affect ATG-9 and a *bona fide* synaptic vesicle protein differently.

We next examined if ATG-9 was abnormally localized to immature autophagosomes. Simultaneous imaging of ATG-9::mCherry and of the autophagosome marker, GFP::LGG-1/Atg8/GABARAP (Alberti et al., 2010, Manil-Segalen et al., 2014, Wu et al., 2015, Stavoe et al., 2016, Hill et al., 2019) in the *unc-26(s1710)/synaptojanin 1* mutants did not reveal enrichment on the same compartments. However, they sometimes appeared adjacent to each other (Figures 3G-3I, 3M).

Synaptojanin plays conserved roles in clathrin-mediated endocytosis, and in *C. elegans*, *Drosophila* and vertebrates, mutations in synaptojanin result in the accumulation of clathrin-coated, abnormal endocytic intermediates (Harris et al., 2000, Verstreken et al., 2003, Cremona et al., 1999, Kim et al., 2002). We examined the relationship between ATG-9 and clathrin by simultaneously imaging ATG-9::GFP and BFP::CHC-1/Clathrin Heavy Chain in the AIY interneurons of wild-type animals and *unc-26(s1710)/synaptojanin 1* mutant animals. We found that in the *unc-26(s1710)* mutants, CHC-1 localized to abnormal foci at synapses (Figures 3K, S3G) and it colocalized well with ATG-9 (Figures 3J-3M). Based on previous findings (Harris et al., 2000, Cremona et al., 1999, Verstreken et al., 2003), these clathrin-rich foci at the synapses probably represent abnormal endocytic intermediates (Figures 3J-3M).

To sum up, ATG-9 accumulates at presynaptic clathrin-rich structures in *unc-26/synaptojanin 1* mutants.

### In autophagy mutants, ATG-9 accumulates into endocytic intermediates at presynaptic sites

The critical role of ATG-9 in autophagy predicts that disruption of autophagy should impact ATG-9 localization (Reggiori et al., 2004, Sekito et al., 2009, Lu et al., 2011). Thus, we also examined ATG-9 localization in mutants with disrupted autophagy (Figure 4A). In loss-of-function alleles *unc-51(e369)/ATG1*, *epg-9(bp320)/Atg101*, *atg-13(bp414)/epg-1* and *epg-8(bp251)/Atg14* (Crawley et al., 2019, Liang et al., 2012, Huang et al., 2013) that affect early steps of autophagosome initiation, we observed abnormal focal accumulation of ATG-9 at presynaptic sites (Figures 4B-4G). Similar to *unc-26* mutants, the abnormal focal accumulation of ATG-9::GFP in *epg-9(bp414)* autophagy mutants co-localized with clathrin heavy chain BFP::CHC-1 (Figures 4I-4K).

**Fig 4.**
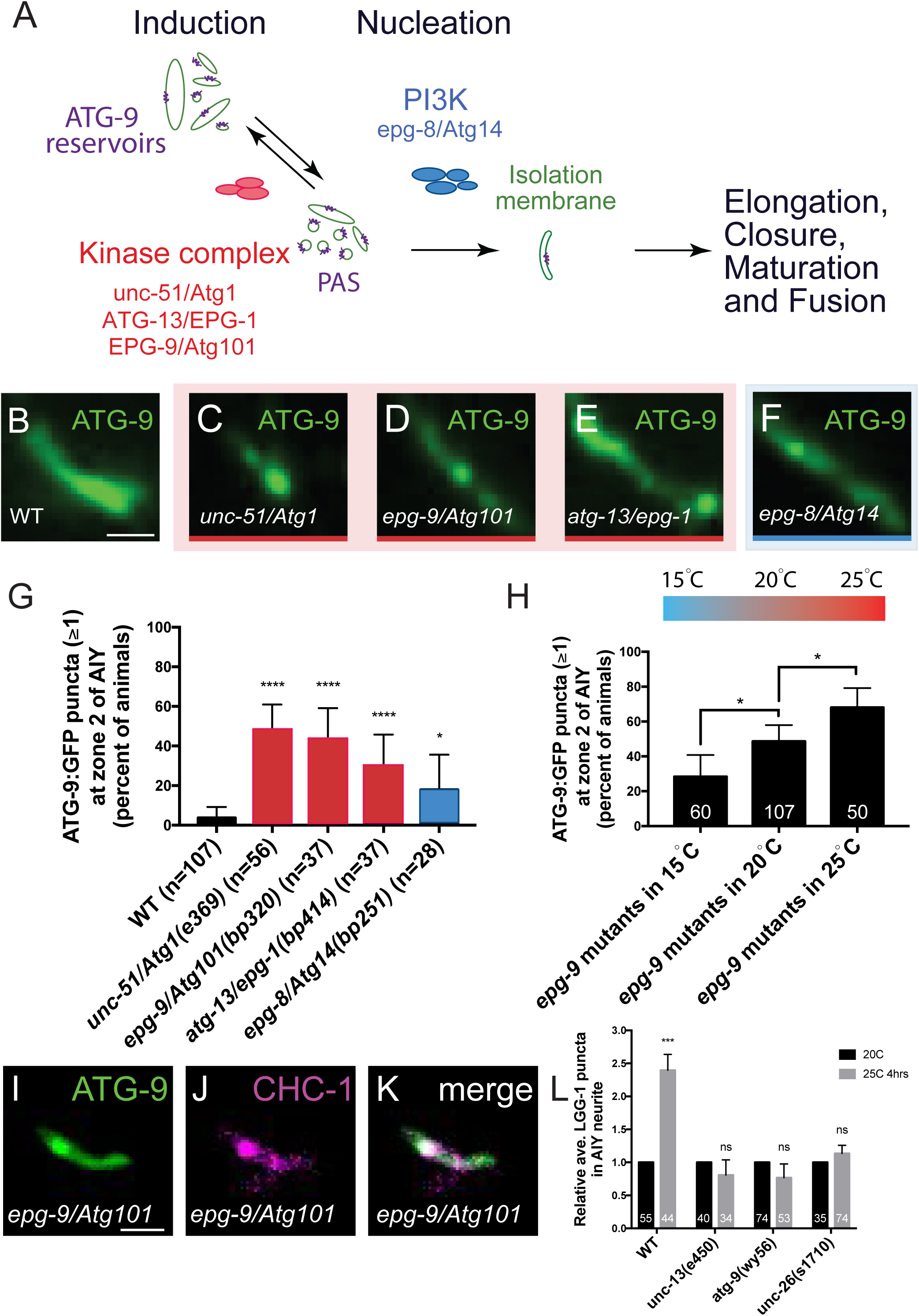
In autophagy mutants, ATG-9 accumulates into endocytic intermediates at presynaptic sites. (A) Schematic of the autophagosome biogenesis pathway. ATG-9 cycles between ATG-9 reservoirs and pre-autophagosomal structures (PAS) during autophagosome biogenesis (Reggiori et al., 2004, Yamamoto et al., 2012, Suzuki et al., 2001). (B-F) Confocal micrographs of ATG-9::GFP at AIY Zone 2 in wild type (B), *unc-51(e369)/Atg1* (C), *epg-9(bp320)/Atg101* (D), *atg-13(bp414)/epg-1* (E), and *epg-8(bp251)/Atg14* (F) mutants. (G) Quantification of percentage of animals displaying one (or more) ATG-9 subsynaptic foci at AIY Zone 2 in wild type and indicated autophagy mutants. The number of animals examined in each condition is indicated by “n”. Error bars show standard error of the mean (SEM). “ns” (not significant), *p<0.05 and ****p<0.0001 (between wild type and the mutants) by two-tailed Fisher’s exact test. (H) Quantification of the percentage of animals with one (or more) ATG-9 subsynaptic foci at AIY Zone 2 in *epg-9(bp320)/Atg101* mutants raised at indicated temperatures. The number of animals examined in each condition is indicated by the numbers on the bars. Error bars show standard error of the mean (SEM). *p<0.01 by two-tailed Fisher’s exact test. (I-K) Representative confocal micrographs of ATG-9::GFP (I), BFP::CHC-1 (pseudo-colored magenta) (J) and merged channels (K) at AIY Zone 2 in *epg-9(bp320)/Atg101* mutants. (L) Quantification of the relative number of LGG-1 puncta in the AIY neurites at 20°C and at 25°C for 4 hours in wild type, *unc-13(e450)*, *atg-9(wy56)* and *unc-26(s*1710) mutants. As primary interneurons in the thermotaxis circuit of *C. elegans*, AIY activity state is found to increase when animals are raised at 25°C for 4 hours, compared with animals at 20°C (Clark et al., 2006, Hawk et al., 2018, Biron et al., 2006). For every genotype, the average number of LGG-1 puncta at 25°C was normalized to the observed average at 20°C to visualize the difference between different neuronal activity state in each genotype. The average number before normalization can be seen in Supplementary Figure 4. “ns” (not significant) and ***p<0.001 (between 20°C and 25°C in each genotype) by Welch’s t test. Scale bars 2μm in (B) for (B)-(F); 2μm in (I) for (I)-(K).

Is the abnormal localization of ATG-9 in early autophagy mutants affected under conditions of increased synaptic activity state and autophagy? We examined ATG-9 in *epg-9(bp414)* mutants reared at 15°C, 20°C and 25°C, and found that the penetrance of the ATG-9 phenotype in the AIY interneurons of *epg-9(bp414)* mutants varied according to the cultivation temperature of the animals (which relates to the activity state of the AIY interneuron (Clark et al., 2006, Hawk et al., 2018); (Figure 4H)). The effects of temperature on the abnormal localization of ATG-9 to foci in autophagy mutants are similar to those observed for ATG-9 in endocytic mutants (Figure 4H and Figure 2N).

To then relate changes in ATG-9 localization at the synapse with activity-dependent increases in synaptic autophagy, we examined LGG-1/Atg8/GABARAP puncta in mutant backgrounds that affect exo-endocytosis at the synapse. Consistent with the previous findings (Hill et al., 2019, Hawk et al., 2018), we observed the average number of LGG-1 puncta increased when the wild type animals were cultivated at 25°C, a condition known to increase the activity state of the AIY neurons (Hawk et al., 2018) and synaptic autophagy (Hill et al., 2019) (Figure 4L, S4A-S4C). Inhibiting exocytosis (in *unc-13(s69)* mutants) or disrupting the autophagy pathway (in *atg-9(wy56)* mutants) eliminated the capacity of the neuron to increase synaptic autophagy in response to increases in the cultivation temperature (Figure 4L, S4C). We observed higher numbers of LGG-1 puncta under basal conditions in *unc-26(s1710)* mutants (Figures S4C). The LGG-1 puncta in *unc-26(s1710)* mutants could not be suppressed by autophagy mutants, suggesting the increased LGG-1 puncta were not bona-fide functional autophagosomes (Figures S4C). Moreover, the LGG-1 puncta in *unc-26(s1710)* mutants did not increase in response to cultivation temperatures that increase the activity state of the neuron (Figures 4L, S4C). Our findings are consistent with previous studies that provide evidence that synaptojanin 1 is important for autophagosome formation when inducing autophagy (Vanhauwaert et al., 2017). We extend these studies, now demonstrating a link between exo-endocytic traffic of ATG-9 at presynaptic sites, and activity-dependent presynaptic autophagy.

### Abnormal accumulation of ATG9A in nerve terminals of mammalian neurons with mutations in endocytic proteins

We next investigated whether the effect of perturbation of endocytosis on ATG9A dynamics in nerve terminals is conserved at mammalian synapses. To this end, we explored the localization of ATG9A in nerve terminals of hippocampal neuronal cultures of mice double KO for dynamin 1 and 3 (the two neuronally enriched dynamin isoforms) and KO for synaptojanin 1 (SJ1) (the neuronally enriched synaptojanin isoform) (Raimondi et al., 2011, Cremona et al., 1999, Ferguson and De Camilli, 2012). Synapses of both genotypes are characterized by a massive accumulation of synaptic vesicle endocytic intermediates, endocytic pits in the case of dynamin mutants and clathrin coated vesicles in the case of synaptojanin 1 mutants. This accumulation is reflected in a very robust clustering in presynaptic terminals of immunoreactive signal for endocytic factors, including clathrin, clathrin adaptors and their accessory proteins such as amphiphysin 2 (Raimondi et al., 2011, Hayashi et al., 2008, Ferguson et al., 2007, Milosevic et al., 2011). Accordingly, anti-amphiphysin 2 immunofluorescence revealed a much stronger synaptic staining in dynamin 1 and 3 DKO neurons, and in SJ1 KO neurons, than in controls (Figures 5). Importantly, anti-ATG9A immunofluorescence also revealed a striking accumulation of this protein at a subset of mutant synapses relative to WT synapses (Figures 5A-5B, 5E-5F). Such hot spots of ATG9A colocalized with amphiphysin 2 immunoreactivity, confirming the synaptic localization of anti-ATG9A (Figure 5). However, the number of synaptic puncta were more numerous for amphiphysin 2 than for ATG9A, suggesting a heterogeneous localization of ATG9A at synapses, or a different impact of the perturbation of dynamin and SJ1 on ATG9A in different neurons.

**Fig 5.**
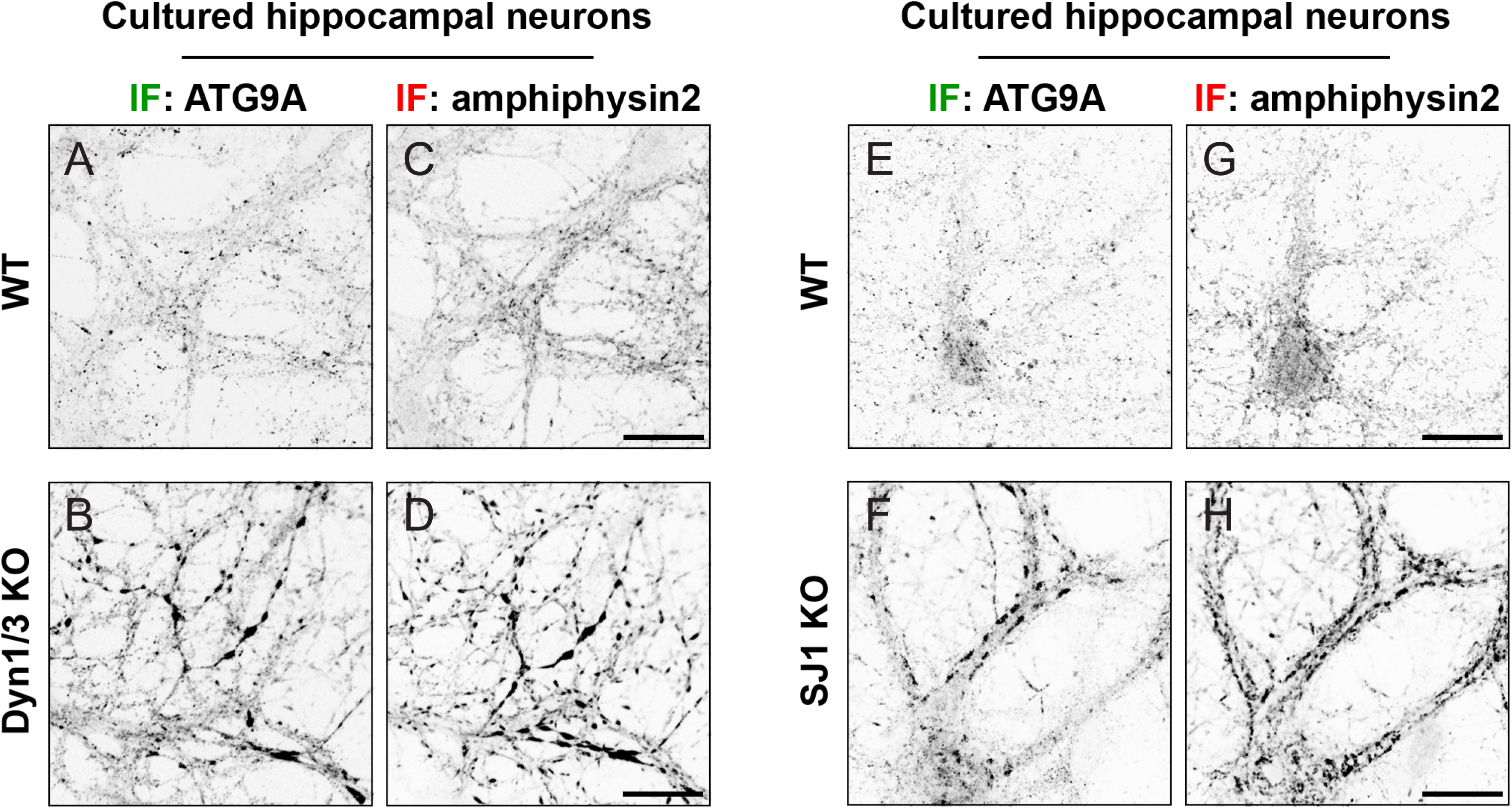
Abnormal accumulation of ATG9A in nerve terminals of mammalian neurons with mutations in endocytic proteins. (A-D) Representative images showing immunoreactivity for ATG9A (A and B) and amphiphysin2 (C and D) in DIV17 hippocampal neuronal cultures from wild type (WT) (A and C) and dynamin 1/3 double KO (Dyn1/3 KO) (B and D) newborn mice. (E-H) Representative images showing immunoreactivity for ATG9A (E and F) and amphiphysin2 (G and H) in DIV23 hippocampal neuronal cultures from wild type (WT) (E and G) and synaptojanin1 KO (SJ1 KO) (F and H) newborn mice. Scale bars 20μm in (C), (D), (G), (H) for (A)-(H).

### A mutation in unc-26/synaptojanin 1 associated with early-onset Parkinsonism (EOP) leads to abnormal focal accumulation of ATG-9 in presynaptic nerve terminals

ATG-9 links autophagy, endocytosis and neuronal activity at synapses. Abnormal function of these processes, which are crucial for maintaining neuronal health and homeostasis, have been implicated in Parkinson’s disease (Vidyadhara et al., 2019, Anglade et al., 1997, Lynch-Day et al., 2012). A missense mutation at an evolutionarily conserved position in the PI4P phosphatase domain of Sac1 domain of SJ1 (R258Q) is associated with early-onset Parkinsonism (Quadri et al., 2013, Krebs et al., 2013). Introduction of the same mutation in mouse and *Drosophila* was reported to affect endocytic trafficking and autophagosome maturation at synapses (Cao et al., 2017, Vanhauwaert et al., 2017). The mutant position, which impairs the catalytic activity of the Sac1 domain, is conserved in *C. elegans* (R216Q) (Figure 6A). Does this mutation also affect ATG9A localization at synapses? We addressed this question both in neuronal cultures of previously described homozygous knock-in mice with the PD mutation (SJ1^RQ^KI mice) (Cao et al., 2017) and in *C. elegans* in which we engineered the homozygous Parkinson’s disease mutation (R216Q) via CRISPR-Cas9.

**Fig 6.**
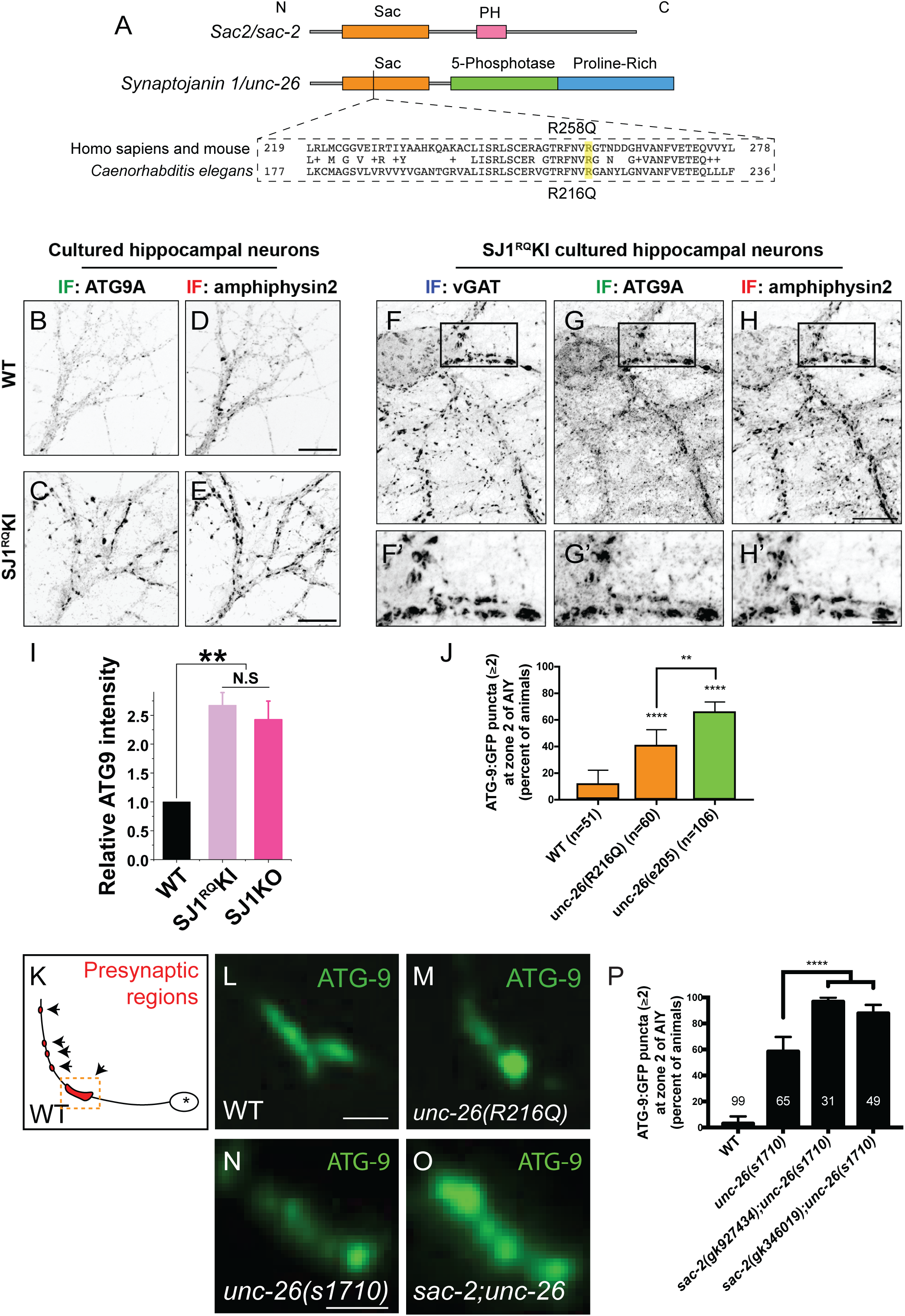
A mutation in unc-26/synaptojanin 1 associated with early-onset Parkinsonism (EOP) leads to abnormal focal accumulation of ATG-9 in presynaptic nerve terminals. (A) Domain structures of *Sac2/sac-2* and *Synaptojanin 1/unc-26*. The mutated residue associated with EOP is conserved (highlighted in yellow). (B-E) Representative images showing immunoreactivity for ATG9A (B and C) and amphiphysin2 (D and E) in DIV23 hippocampal neuronal cultures from newborn wild type (WT) mice (B and D) and mice harboring a EOP associated mutation in the synaptojanin1 (SJ1^RQ^KI) gene (C and E). (F-H, F’-H’) Representative images showing immunoreactivity for vesicular GABA transporter (vGAT) (F), ATG9A (G) and amphiphysin2 (H) in DIV19 hippocampal neuronal cultures from SJ1^RQ^KI newborn mice. (F’-H’) enlarged images of squared regions in (F-H). (I) Quantification of relative ATG9A intensity in nerve terminals of wild type (WT), SJ1^RQ^KI and SJ1KO hippocampal neuronal cultures. n = 3 independent cultures. Error bars show standard error of the mean (SEM). **p<0.01 by Student’s t test. (J) Quantification of abnormal ATG9 localization in *unc-26(R216Q)* and *unc-26(e205)* worm mutants. The bars show *C. elegans* that display two (or more) ATG-9 subsynaptic foci at Zone 2 in wild type. The number of animals examined in each condition is indicated by “n”. Error bars show standard error of the mean (SEM). **p<0.01 and ****p<0.0001 (between wild type and the mutants, and between the two mutant groups) by two-tailed Fisher’s exact test. (K) Schematic of an AIY interneuron with presynaptic sites (in red and arrowheaded) and Zone 2 highlighted by dashed square. (L-O) Representative confocal micrographs of ATG-9::GFP at AIY Zone 2 in wild type (L), *unc-26(R216Q)* (M) mutants, *unc-26(s*1710) (N) and *sac-2(gk346019);unc-26(s1710)* (O) mutants. (P) Quantification of the percentage of animals displaying two (or more) ATG-9 subsynaptic foci at AIY Zone 2 in wild type, *unc-26(s*1710)*, sac-2(gk927434);unc-26(s1710)* and *sac-2(gk346019);unc-26(s1710)* mutants. The number of animals examined in each condition is indicated by the numbers on the bars. Error bars show standard error of the mean (SEM). ****p<0.0001 by two-tailed Fisher’s exact test. Scale bars 20μm in (D) for (B) and (D); 20μm in (E) for (C) and (E); 20μm in (H) for (F)-(H); 5μm in (H’) for (F’)-(H’); 2μm in (L) for (L)-(M); 2μm in (N) for (N)-(O)

Immunofluorescence staining of endogenous ATG9A in hippocampal cultures generated from SJ1^RQ^KI mice revealed abnormal ATG9A accumulations that colocalized with focal enrichment of amphiphysin 2 foci at synapses (Figures 6B-6E, 6I) and were similar to those observed in SJ1 KO neurons (Figures 5E-5H). As in the case of dynamin 1 and 3 DKO synapses and of SJ1 KO synapses (Hayashi et al., 2008, Raimondi et al., 2011), such accumulations were typically more prominent in inhibitory presynaptic GABA-ergic nerve terminals – as revealed by immunostaining for vGAT (Figures 6F-6H, 6F’-6H’), which generally have higher levels of tonic activity.

Likewise, in *unc-26 (R216Q)* mutant *C. elegans*, ATG-9::GFP was abnormally localized to subsynaptic foci which resembled those observed in *unc-26(e205)/SJ1* null alleles - and accordingly were enriched in clathrin - albeit with a lower penetrance, consistent with partial loss of function (Figures 6J-6M, S5H-S5J).

We also observed that the *unc-26(R216Q)* allele, contrary to the loss-of-function allele, did not produce obvious changes in the localization of the synaptic vesicle proteins SNG-1 or RAB-3 at presynaptic regions (Figures S5A-S5G), showing that *unc-26 (R216Q)* differentially affects ATG-9 localization and synaptic vesicle protein localization. These findings are consistent with *Drosophila* studies indicating that the *unc-26(R216Q)* Parkinson’s disease mutation impairs autophagy at the synapse (Vanhauwaert et al., 2017), and are extended to show an impact of the mutation on ATG-9 trafficking at presynaptic sites.

Another Sac domain-containing protein, Sac2/INPP5F (Fig. 6A), is located within a Parkinson’s disease risk locus identified by genome-wide association studies (Blauwendraat et al., 2019, Nalls et al., 2014). We examined two putative null alleles of SAC-2 in *C. elegans*, *sac-2(gk927434)* and *sac-2(gk346019)*, for phenotypes in ATG-9 localization. While single mutants of *sac-2* do not affect ATG-9 localization (data not shown), *sac-2(gk927434);unc-26(s1710)* and *sac-2(gk346019);unc-26(s1710)* double mutants enhance the abnormal localization of ATG-9 in *unc-26(s1710)* single null allele, suggesting that *sac-2* and *unc-26/synaptojanin 1* function synergistically in mediating ATG-9 trafficking at synapses (Figures 6N-6P). Our observations are consistent with previous findings showing that Sac2 and synaptojanin 1 have overlapping roles in the endocytic pathway at synapses (Cao et al., 2020). Importantly, our findings indicate that lesions associated with early onset Parkinsonism in endocytic mutants result in abnormal ATG-9 accumulation, suggesting a possible link between this condition, ATG-9 traffic at synapses and autophagy.

## Discussion

In *C. elegans*, ATG-9 exits the Golgi complex in an AP-3 dependent manner. We had previously shown that ATG-9, the only intrinsic transmembrane protein of the autophagy machinery, is transported by UNC-104/KIF1A to presynaptic nerve terminals, where it plays a critical role for synaptic autophagosome formation (Stavoe et al., 2016). Here we show that AP-3 is a critical component of the coat that sorts ATG-9 in the vesicles targeted to presynaptic sites in *C. elegans*. The *C. elegans* AP-3 protein complex is structurally and mechanistically related to the mammalian AP-4 complex, which in vertebrates is required for signal-mediated transport of ATG9 from the TGN to the peripheral cytoplasm (Rout and Field, 2017, Dell’Angelica, 2009). Invertebrates, including *C. elegans*, do not have AP-4 complexes. Whether AP-3 has a role for ATG-9 export from the Golgi complex in mammalian cells remains to be explored.

ATG-9 enriched vesicles at the synapse might represent a distinct subpopulation of vesicles. In nerve terminals, as shown by our EM analyses, the bulk of ATG-9 is localized in small vesicles. Interestingly, while one cannot identify molecularly distinct vesicle populations based on size and morphological appearance in EM, immunogold staining suggests a predominant concentration of ATG-9 on a subpopulation of vesicles. Likewise, fluorescent microscopy revealed only partial overlap in nerve terminals between the distribution of ATG-9 and of the intrinsic membrane protein of synaptic vesicles, SNG-1/synaptogyrin. Consistent with these findings, ATG-9 was identified by mass spectrometry in a synaptic vesicle fraction obtained by immunopurification of vesicles positive for the synaptic vesicle protein synaptophysin or by differential and Ficoll density gradient centrifugation (Boyken et al., 2013, Chantranupong et al., 2020), suggesting compositional overlap between ATG-9 vesicles and bona fide synaptic vesicles.

Synaptically-localized ATG-9 positive vesicles undergo exo-endocytosis. We demonstrate that their exocytosis is *unc-13/unc-18*-dependent, and that their endocytosis is affected by *dyn-1/dynamin*, *unc-26/synaptojanin, unc-57/endophilin* and *unc-11/AP180*, all genes required for synaptic vesicle endocytosis. In *unc-26* mutants, ATG-9 predominantly accumulates in foci which are also enriched in clathrin. The observation that loss-of-function mutations of *unc-13/unc-18* suppress the abnormal distribution of ATG-9 in endocytic mutants shows that such redistribution is the result of abnormal endocytosis after exocytosis. Consistent with these findings in *C. elegans*, in mice a robust accumulation of ATG-9 was detected in a subpopulation of neurons that harbor loss-of-function mutations in the genes that encode neuronal isoforms of dynamin and synaptojanin. These findings are also consistent with studies in non-neuronal mammalian cells showing an accumulation of ATG9 at the plasma membrane upon perturbation of dynamin-dependent endocytosis, as detected by fluorescence microscopy (Puri et al., 2013, Puri et al., 2014, Popovic and Dikic, 2014) or a cell surface biotinylation assay (this study).

The exo-endocytosis of ATG-9 at synapses reveals a link between synaptic vesicle traffic and autophagy. We observe that disruptions in the synaptic vesicle cycle, or autophagy, similarly result in abnormal accumulation of ATG-9 at the plasma membrane and in clathrin-rich presynaptic foci. We interpret these clathrin-rich foci to be endocytic intermediates onto which ATG-9 gets trapped in the endocytic and autophagy mutants.

A missense mutation in the endocytic protein synaptojanin in *C. elegans* (corresponding to human R258Q associated with early-onset parkinsonism (EOP)) results in abnormal accumulation of ATG-9 in clathrin-rich synaptic foci. Synaptojanin contains two phosphatase domains: an inositol 5-phosphatase domain which has been associated with most of the roles of synaptojanin in the endocytic trafficking of synaptic vesicles, and an inositol 4-phosphate Sac1 phosphatase domain, which can dephosphorylate to some extent also PI3P and PI(3,5)P2, and whose precise physiological function is less understood. The R258Q mutation, which selectively abolishes the activity of the Sac1 phosphatase domain, was shown to impair presynaptic endocytic flow, more prominently at inhibitory synapses, which have generally higher tonic activity (Cao et al., 2017), and also impair autophagy (Vanhauwaert et al., 2017). Our findings are consistent with an impact of the EOP mutation on autophagy, as we demonstrate that ATG-9 is mislocalized at synapses both in *C. elegans* harboring the homologous *unc-26(R216Q)* lesion and in the R258Q mutant mice. In *Drosophila*, the corresponding EOP mutation in synaptojanin also resulted in neurodegeneration (Vanhauwaert et al., 2017). In view of the role of autophagy in the control of nerve terminal health and homeostasis, the defect in autophagy may contribute to the neurodegeneration leading to EOP.

Together, our data support a model whereby ATG-9 couples the synaptic exo-endocytosis and autophagy (Figure 7). ATG-9 is critical for autophagosome biogenesis, and by trafficking via exo-endocytosis at presynaptic sites, ATG-9 could coordinate synaptic autophagy with synaptic vesicle recycling, linking synaptic autophagy to the activity state of the neuron.

**Fig 7.**
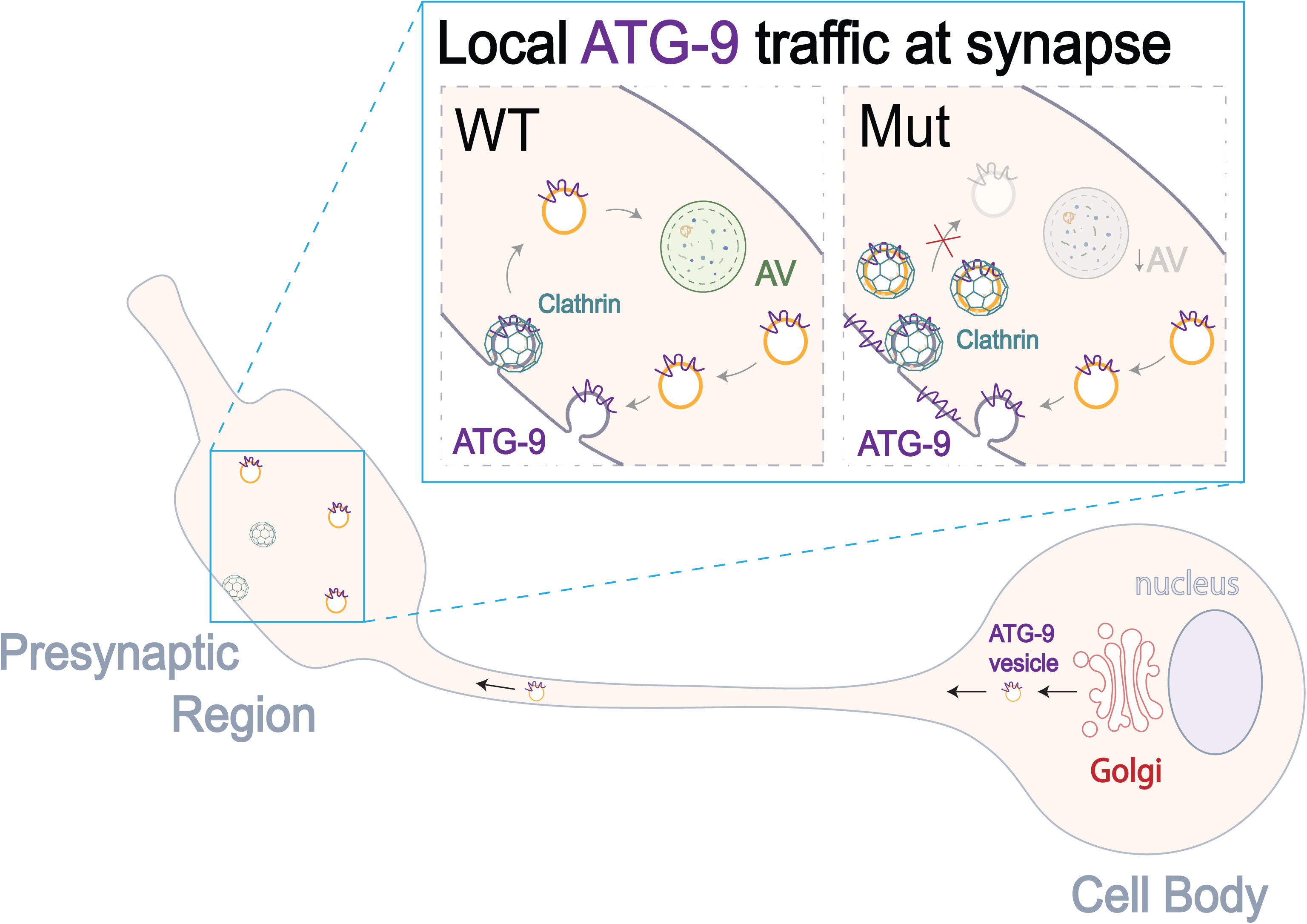
ATG-9 traffic in neurons: a model from origin in the cell body to local traffic at the synapse. Schematic model of ATG-9 trafficking in *C. elegans* neurons. ATG-9 vesicles originate from the trans-Golgi network via AP-3-dependent budding (*C. elegans* lacks distinct AP-3 and AP-4 adaptors). ATG-9 vesicles undergo anterograde transport to the synapses, which depends on the kinesin UNC-104/KIF1A (Stavoe et al., 2016). Once ATG-9 vesicles reach the presynaptic region, they undergo exo-endocytosis. In mutants that disrupt endocytosis, or in autophagy mutants, ATG-9 accumulates into clathrin-enriched synaptic foci, and activity-dependent presynaptic autophagy is compromised. Similar mechanisms operate at mammalian nerve terminals.

## Acknowledgments

We thank Erik Jorgensen (Department of Biology, the University of Utah), Kang Shen (Department of Biology, Stanford University) and Hong Zhang (Department of Cell Biology, University of Massachusetts Medical School) for providing strains and constructs. We thank Center for Cellular and Molecular Imaging, Electron Microscopy Facility at Yale Medical School for assistance with the work presented here, and Yumei Wu, Irina Kolotuev, David Hall, Maike Kittelmann and Szi-chieh Yu for advice on immunoelectron microscopy experiments. We thank current and past members of the Colón-Ramos lab for help, advice and insightful comments on the project. We thank the Caenorhabditis Genetics Center (funded by NIH Office of Research Infrastructure Programs P40 OD010440) for *C. elegans* strains. S.Y. was supported by China Scholarship Council-Yale World Scholars Program. I.O. summer research was supported by Howard Hughes Medical Institute Exceptional Research Opportunities Program (HHMI ExROP) and Yale BioMed Amgen Scholars Program. Research in the P.D.C. lab was supported by the NIH (NS36251 and DA18343), the Parkinson’s Foundation, the Kavli Foundation, MJFF (ASAP-000580) and a fellowship from the National Research Foundation of Korea to D.P. (2019R1A6A3A03031300). Research in the D.A.C.-R. lab was supported by NIH R01NS076558, DP1NS111778 and by an HHMI Scholar Award.

## Author contributions

Conceptualization: S.Y., S.H. and D.A.C.-R.; methodology: S.Y., L.M., S.H., M.C., L.S. and D.A.C.-R.; software: L.S.; investigation in *C. elegans*: S.Y., L.M., S.H., Z.X., I.G. and I.O.; investigation in mammalian cells: D.P. and M.C.; formal analysis: S.Y., L.M. and D.P.; writing - original draft: S.Y., L.M., P.D.C. and D.A.C.- R.; writing - review and editing: S.Y., L.M., S.H., Z.X., L.S., D.P., M.C., P.D.C. and D.A.C.-R.; visualization: S.Y., L.M., D.P. and D.A.C.-R.; supervision, project administration and funding acquisition: P.D.C. and D.A.C.-R..

## Figure Legends

**Fig S1, related to Fig 1. ATG-9 localizes to a subset of vesicles at synapses and does not extensively colocalize with ER or mitochondrial markers at cell body; RAB-3 is not affected by APB-3.**

(A-B) Immunogold electron microscopy of nerve terminals transgenic worms expressing ATG-9::GFP, and done with antibodies directed against GFP. Note that immunogold particles are enriched at specific presynaptic areas. The similar localization of immunoreactivity in closely adjacent sections (A) and (B) speaks against the possibility that the non-homogenous distribution of gold particles in the terminal may be a labeling artifact. Blue line, outline of the nerve terminals. Dense projections are shaded in orange. “m”, mitochondria.

(C) Quantification of the percentage of immunogold particles on indicated subcellular structures (vesicles, plasma membrane or mitochondria, reported as the percent of total particles in the portion of the neurite visible in the section).

(D) Quantification of the number of immunogold particles on subcellular structures (vesicles, plasma membrane or mitochondria) divided by area occupied by the cellular structures (μm^2^).

(E) Schematic of a cell body and organelles in the AIY interneurons.

(F-H) Confocal micrographs of SP12::GFP (ER marker) (F), ATG-9::mCherry (G) and merged channels (H) at the cell body of AIY.

(I-K) Confocal micrographs of TOMM-20::GFP (mitochondrial marker) (I), ATG-9::mCherry (J) and merged channels (K) at cell body of AIY.

(L) Schematic of an AIY interneuron with RAB-3 localization (red), and synaptic-rich region of Zone 2 in orange dashed box.

(M-O) Confocal micrographs of RAB-3::mCherry at Zone 2 in wild type (M) and *apb-3(ok429)* mutants (N), and quantification (O). Error bars show standard error of the mean (SEM). “ns”: not significant by Welch’s t test between wild-type and mutant animals. Each dot in the scatter plot represents a single animal.

Scale bars 200nm in (A) for (A)-(B) and in inset; 5μm in (F) for (F)-(K); 2μm in (M) for (M)-(N)

**Fig S2, related to Fig 2. Endocytosis regulates ATG-9 localization at synapses.**

(A) Quantification of the percentage of animals displaying two (or more) ATG-9 foci at AIY Zone 2 synapses in wild type and endo-exocytic mutants. The number of animals examined in each condition is indicated by “n”. Error bars show standard error of the mean (SEM). ***p<0.001 and ****p<0.0001 by two-tailed Fisher’s exact test.

(B) Mean intensity of ATG-9 at AIY Zone 2 in wild type and *unc-26(s1710)* mutants. Error bars show standard error of the mean (SEM). *p<0.05 (between wild type and the mutants) by Welch’s t test. Each dot in the scatter plot represents a single animal.

(C-D) Confocal micrographs of ATG-9::GFP in the AIY Zone 2 for *unc-26(e205)*

(C) mutants, *unc-26(e205)* mutants with UNC-26 cDNA rescue array cell-specifically expressed in AIY (D).

(E) Quantification of the percentage of animals displaying one (or more) ATG-9 foci at the synapse-rich AIY Zone 2 in wild type, *unc-26(s1710)* mutants and *unc-26(s1710)* mutants with an UNC-26 cDNA rescue array cell-specifically expressed in AIY. The number of animals examined in each condition is indicated by the “n”. Error bars show standard error of the mean (SEM). “ns” (not significant), ****p<0.0001 by two-tailed Fisher’s exact test.

(F-G) Confocal micrographs of ATG-9::GFP in the AIY Zone 2 in *unc-13(s69)* (F) and *unc-18(e81)* (G) mutants.

Scale bars 2μm in (C) for (C)-(D); 2μm in (F) for (F)-(G)

**Fig S3, related to Fig 3. Defects of synaptic vesicle clusters and clathrin heavy chain (CHC-1) in *unc-26* mutants as compared to wild type.**

(A) Immunoblots (IB) for dynamins in the Dyn1/2 CKO fibroblast, and controls.

(B) Quantification of the dynamin levels in Dyn1/2 CKO fibroblasts relative to a tubulin control, in Dyn1/2 CKO and the control cells (CTL). n = 3 independent experiments. Error bars show standard error of the mean (SEM). **p<0.01 by Student’s t test.

(C-E) Confocal micrographs of RAB-3::mCherry (pseudo-colored green) in wild type (C) and *unc-26(s1710)* mutants (D), and SNG-1::GFP (E) in the *unc-26(s1710)* mutants. Note that like ATG-9, synaptic vesicle associated protein RAB-3 concentrates near synaptic sites in wild type animals (arrows, compare to Figure 1), but unlike ATG-9, these synaptic vesicle proteins become diffusely localized in *unc-26(s1710)* mutants (consistent with observations from (Harris et al., 2000, Verstreken et al., 2003, Ferguson et al., 2007, Raimondi et al., 2011, Milosevic et al., 2011)).

(F) Quantification of the distribution of RAB-3 in wild type and *unc-26(s1710)* mutants for the synaptic region in C and D, and as described (STAR Methods). **p<0.01 (between wild type and the mutants) by Welch’s t test. Each dot in the scatter plot represents a single animal.

(G) Quantification of CHC-1 clusters intensity at synapses per AIY neuron in wild type and *unc-26 (s1710)* mutants, and as described (STAR Methods). *p<0.05 (between wild type and the mutants) by Welch’s t test. Each dot in the scatter plot represents a single animal. The method described in STAR Methods.

**Fig S4, related to Fig 4. Activity-dependent autophagy at presynaptic sites.**

(A-B) Confocal micrographs of cytoplasmic mCherry and eGFP::LGG-1 at AIY Zone 2 in wild type (A) and *unc-26(s1710)* mutants (B). Yellow dashed lines in AIY emphasize the synaptic region, with the yellow dashed box emphasizing the synaptic-rich AIY Zone 2 region of AIY, and the white dashed line, the asyntaptic region. The arrow points to autophagosomes (as visualized with eGFP::LGG-1 (Alberti et al., 2010, Manil-Segalen et al., 2014, Wu et al., 2015, Stavoe et al., 2016, Hill et al., 2019)).

(C) Quantification of the average number of LGG-1 puncta in the AIY neurites at 20°C and at 25°C for 4 hours in wild type, *unc-13(e450)*, *atg-9(wy56), unc-26(s1710)* and *unc-26(s1710);atg-9(wy56)* mutants. The number of animals examined in each condition is indicated by the numbers on the bars. *p<0.05 and ****p<0.0001 by ordinary one-way ANOVA with Dunnett’s multiple comparisons test between wild-type and the mutant groups.

Scale bars 5μm in (A) for (A)-(B).

**Fig S5, related to Fig 6. The lesion associated with EOP does not affect localization of synaptic vesicle proteins.**

(A) Schematic of an AIY interneuron.

(B) Quantification of the distribution of SNG-1 at Zone 3 in wild type and *unc-26(R216Q)* mutants as described (STAR Methods). Error bars show standard error of the mean (SEM). “ns” (not significant) (between wild type and the mutants) by Welch’s t test. Each dot in the scatter plot represents a single animal.

(C-G) Representative confocal micrographs of RAB-3::mCherry at Zone 3 in wild type (C), *unc-26(R216Q)* (D), *unc-26(s1710)* (E) mutants; SNG-1::GFP in wild type

(F) and *unc-26(R216Q)* (G) mutants. The arrowheads denote the clusters at Zone 3. The brackets denote the diffuse RAB-3 localization at Zone 3.

(H-J) Representative confocal micrographs of BFP::CHC-1 (pseudo-colored magenta) (H), ATG-9::GFP (I) and merged channels (J) at Zone 2 in *unc-26(R216Q)* mutants.

Scale bars 5μm in (C) for (C)-(G); 2μm in (H) for (H)-(J);

## STAR Methods

### CONTACT INFORMATION

Further information and requests for resources and reagents should be directed to and will be fulfilled by the Lead Contact, Daniel A. Colón-Ramos.

### EXPERIMENTAL MODEL AND SUBJECT DETAILS

#### *C. elegans* Strains and Genetics

Worms were raised on nematode growth media plates at 20°C or room temperature using OP50 *Escherichia coli* as a food source (Brenner, 1974). Animals were analyzed as larva 4 (L4) stage hermaphrodites. For wild-type nematodes, *C. elegans* Bristol strain N2 was used. For a full list of strains used in the study, please see the KEY RESOURCES TABLE.

#### Molecular Biology and Transgenic Lines

*C. elegans* transgenic strains were created by injecting pSM vector-derived plasmids (listed on KEY RESOURCES TABLE) by standard injection techniques with co-injection markers punc-122::gfp and punc-122:rfp.

The cDNA constructs (UNC-26, TGN-38, clathrin heavy chain (CHC-1)) generated were amplified from a cDNA pool of a mixed population of *C. elegans*. Detailed sub-cloning information is available upon request.

### METHOD DETAILS

#### *C. elegans* CRISPR Transgenics

To introduce the lesion in unc-26/synaptojanin 1 associated with early-onset Parkinsonism (EOP), we used CRISPR-Cas9 to substitute CGA with CAA, and generate the homozygous mutation R216Q. CRISPR transgenic strain was generated by precision genome editing method using CRISPR-Cas9 and linear repair templates, as previously described (Paix et al., 2017) and using the targeted gene crRNA (GGCACUCGAUUCAACGUAC) and repair template ssODN (GACGTGTTGCTCTAATATCTCGTCTAAGTTGTGAGCGTGTCGGCACTCGATTCAACGTACAAGGAGCCAATTATCTCGGAAATGTGGCTAATTTCGTCGAGACTGAGCAATTGTTGCTTTT)

#### Cell Autonomy and Rescue of *unc-26*

The *unc-26* mutant phenotype was rescued by cell specifically expressing the wild type *unc-26* cDNA under AIY-specific *ttx-3* promoter fragment (Colon-Ramos et al., 2007).

#### Fluorescence Microscopy and Confocal Imaging

We imaged *C. elegans* by using 60x CFI Plan Apo VC, NA 1.4, oil objectives on an UltraView VoX spinning disc confocal microscope and on a NikonTi-E stand (PerkinElmer) with a Hammamatsu C9100-50 camera. *C. elegans* were immobilized on a 2% agarose pad with 10mM levamisole (Sigma). Images were processed with Volocity software or Fiji.

#### Inhibiting Oxidative Phosphorylation Using a microfluidic-hydrogel device

To inhibit oxidative phosphorylation by hypoxia, a reusable microfluidic polydimethylsiloxane (PDMS) microfluidic device was used, as described (Jang et al., 2020) while imaging ATG-9::GFP localization in *pfk-1.1(gk922689);olaIs34* (pttx-3::atg-9::gfp and pttx-3::mCherry::rab-3). Normoxic and hypoxic conditions were applied to animals for 10 min sequentially by alternating the flow of air and nitrogen gas, respectively. As a positive control, synaptic vesicle protein RAB-3 was also imaged as reported (Jang et al., 2016) (Figure 2M).

#### Electron Microscopy

Worms were prepared by high pressure freeze and freeze substitution as described (Rostaing et al., 2004, Manning and Richmond, 2015, Kolotuev et al., 2012). Briefly, ∼10-20 worms at the L4 stage were loaded into carriers coated lightly with hexadecane (Specimen carrier Type A and B, Technotrade International) with 20% BSA and *E. coli* for high-pressure freezing (Leica EM HPM 010). After freezing, samples were transferred to an AFS machine for freeze substitution (Leica EM AFS2) using a custom made workbox submerged in liquid nitrogen. Samples were incubated in .1% uranyl acetate in anhydrous acetone as follows: −90C for 48 hours, temperature raised to −50C over 8 hours, held at −50C for 12 hours. Next, samples were washed with ethanol several times over 2 hours and incubated in graded concentrations of HM20 resin (Lowicryl HM20, Electron Microscopy Sciences) in ethanol (3 hours at 25% HM20 in EtOH, 3 hours in 50%, 16h in 75%, 6 hours in 100% HM20). Finally, worms were embedded in HM20 resin at −50C within the AFS chamber. To facilitate embedding, we used a custom-made aluminum chamber similar to that described in (Kolotuev, 2014). Carriers containing the worms were inverted onto a small square of Aclar (Sigma) and manipulated using fine needle tips to dissociate worms from the planchette. Worms were then placed onto a small drop of HM20 within a double-sided adhesive frames (Thermo Scientific) sandwiched between squares of Aclar. Thin embedding at this step was critical to see the transparent worms later during sectioning. Samples were cured under UV light for 48 hours at −50C, brought to room temperature over 14 hours, and remained under UV light for another 24 hours at room temperature.

To facilitate sectioning, fixed worms were cut from the embedded square and glued onto thin plastic blocks made using Epon resin in a Chang mold (EMS). Worms were trimmed (Diatome Trim 45) and sectioned (Diatome 4.0 Ultra) on a Leica UC7 (Leica) until the desired area was identified. 500 nm thick sections were collected and stained with toluidine blue to check for the desired ROI.

The nerve ring and AIY Zone 2 were identified using anatomical landmarks described in the original *C. elegans* connectome (White et al., 1986). The nerve ring is located in the head of the animal and forms a ring of densely packed neurites around the pharynx. AIY Zone 2 synapses are positioned in a ventral bundle of neurites just posterior to the nerve ring. These synapses reside at the ventral base of the neurite bundle and form a unique humped shape with multiple dense projections. The left and right process of AIY contact one another at the posterior end of Zone 2 synapses. Chemical synapses in *C. elegans* are defined by the presence of presynaptic dense projection in the neurite.

50 nm thick sections were collected from at least one animal per genotype on nickel slot grids covered with Formvar (EMS). When possible, serial sections were collected. Antibody staining was performed within one day. Grids were incubated for 10 minutes in .15% glycine and .1M ammonium chloride in PBS, followed by incubation in blocking solution (1% BSA and 1% CWFS gelatin in PBS) for 10 minutes. Grids were then incubated in anti-GFP primary antibody diluted in 2 parts PBS and 1 part blocking solution overnight at 4C (ab6556 1:20, Abcam) After washing in PBS four times over 15 minutes, grids were incubated in Protein A Gold conjugated to 10 nm particles diluted in 2 parts PBS and 1 part blocking solution (1:75, University Medical Center Utrecht) for 30 minutes. Grids were washed again in PBS four times over 15 minutes, followed by 5 minutes incubation in 1% glutaraldehyde in PBS (EMS), and three quick washes on water. After drying, grids were post stained with Reynold’s lead citrate for 4 minutes, 2.5% uranyl acetate for 4 minutes, and lead citrate for 1 minute and allowed to dry at least 1 hour before imaging.

Images were acquired on a FEI Tecnai Biotwin (FEI) equipped with a SIS Morada 11 megapixel CCD camera and a TALOS L120 (Thermo Fisher) equipped with a Ceta 4k x 4k CMOS camera. For serial sections, images were aligned in z using the TrakEM2 plugin in FIJI.

#### Hippocampal Neuron Culture

Mice were maintained on the C57BL6/129 hybrid genetic background. Heterozygous mice were mated to generate homozygous knockout or knock-in with their littermate controls. For neuronal cultures, P0 pups were genotyped by PCR and then hippocampi were dissected. Tissues were digested for 20 min in a papain/HBSS solution (20 U/ml) containing DNase (20 μg/ml). Cells were dissociated by trituration and then plated onto poly-d-lysine coated coverslips. After 3 hours incubation, the plating medium was exchanged to complete neurobasal medium (2% B-27 and 0.5 mM L-glutamine in neurobasal medium). Cells were maintained at 37°C in a 5% CO_2_ humidified incubator and the 30% of cultured medium was replaced with new complete neurobasal medium at 4, 7 and 14 days in vitro (DIV). All adult mice for breeding were maintained on a 12 hours light/dark cycle with standard mouse chow and water ad libitum. All research and animal care procedures were approved by the Yale University Institutional Animal Care and Use Committee.

#### Immunofluorescence

Cultured hippocampal neurons were briefly washed in a pre-warmed tyrode (136 mM NaCl, 2.5 mM KCl, 2 mM CaCl_2_, 1.3 mM MgCl_2_, 10 mM HEPES and 10 mM glucose) and then fixed with 4% PFA in 4% sucrose-containing 0.1M PB buffer (pH7.3) for 15 min at RT. After fixation, cells were washed in PBS and incubated with blocking buffer (3% BSA, 0.2% Triton X-100 in PBS) for 30 min at RT. Primary (1 hour) and secondary antibody (45 min) incubations were subsequently performed in the blocking buffer at RT. After washing, samples were mounted on slides with Prolong Gold antifade reagent (Invitrogen).

#### Dynamin Conditional Knockout

For the tamoxifen inducible KO, Dynamin 1/2 conditional knockout (CKO) cells (Ferguson et al., 2009) were treated with 2 μM 4-hydroxy-tamoxifen for 2 days and then further incubated with 300 nM 4-hydroxy-tamoxifen for 3 days. Depletion of total dynamin levels was confirmed by western blotting.

#### Surface Biotinylation

Control and dynamin 1/2 conditional KO cells were washed three times with ice-cold PBS and incubated with ice-cold EZ-Link Sulfo-NHS-LC-Biotin (0.25 mg/ml) in PBS for 30 min at 4 °C to label the surface proteins. Unbound biotins were quenched and removed by 50 mM glycine in ice-cold PBS for 10 min at 4 °C. After washout, cells were lysed with 1% triton X-100 lysis buffer (20 mM Tris-HCl, pH 8, 1% triton X-100, 10% glycerol, 137 mM NaCl, 2 mM EDTA, 1 mM PMSF, 10 mM leupeptin, 1.5 mM pepstatin and 1 mM aprotinin) and centrifuged at 14,000 g for 20 min at 4°C. The supernatants were collected, and protein concentrations were determined by the BCA Protein Assay Kit. Same amount of lysates (600 μg) were incubated with NeutraAvidin particles for 2 hours at 4 °C to pull-down the biotinylated proteins. Particles were washed three times by lysis buffer, eluted with 2x sample buffer and boiling (5 min) and the eluates were processed for SDS-PAGE and western blotting. The level of proteins were quantified by densitometry using Fiji.

### QUANTIFICATION AND STATISTICAL ANALYSIS

#### *C. elegans* AIY Quantifications

##### Presynaptic Enrichment

Morphologically, AIY can be divided into four segments, consistent with the *C. elegans* EM reconstructions (White et al., 1986): the cell body; a proximal asynaptic region termed Zone 1; a ∼5μm synaptic-rich region termed Zone 2, located at the turn of the neuron into the nerve ring; and Zone 3, which is the distal part of the neurite located at the nerve ring.

ATG-9 or RAB-3 enrichment at synapses was quantified in the integrated transgenic line *olaIs34*, expressing pttx-3::atg-9::gfp and pttx-3::mCherry::rab-3 in the wild type and *apb-3(ok429)* mutants. Fluorescence intensity at Zone 2 was quantified in Fiji (Schindelin et al., 2012) in maximal projection confocal micrographs. ATG-9 (or RAB-3) enrichment at synapses represents Zone 2 signal subtracted by average cytoplasmic signal at the cell body (Figures 1X and S1O). Ratio of ATG-9 intensity of cell body/synapses was quantified in *olaIs34* in the wild type, *apb-3(ok429)*, *apm-3(gk771233)* and *apd-3(gk805642)* mutants. In maximal projection confocal micrographs, fluorescence intensities were measured, background-subtracted (from cytoplasmic signal in the cell body) and averaged for different subcellular regions (using an identically-sized oval-shaped object). The ‘Ratio between ATG-9 intensity at the cell body and synapses’ represents signal of ATG-9 intensity (after the processing) at the cell body divided by the intensity (after the processing) at Zone 2 (as shown in the equation below; Figure 1Y).

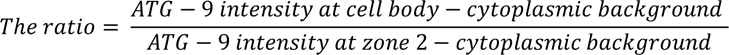

To quantify ratio of SNG-1 intensity of cell body/synapses, we used the extrachromosomal line *olex4060* in the wild type and *apb-3(ok429)* mutants. Maximal projection confocal micrographs were taken on SNG-1 and measured as the intensity at Zone 2 by segmented line scans and the intensity at the cell body by the oval selections of the whole cell body. The ratio of SNG-1 intensity between cell body and synapses was reported as the intensity of SNG-1 at cell body divided by the intensity of SNG-1 at Zone 2 (Figure 1Z).

#### Penetrance of ATG-9:GFP Subsynaptic Foci at the Presynaptic Region

To quantify the penetrance of ATG-9 subsynaptic foci at the presynaptic region (Zone 2), we used integrated transgenic strain *olaIs34* and *olaIs33* in the wild type and mutant background animals. For the temperature sensitive allele *dyn-1(ky51)* and the wild-type animals, animals were held at either 20°C (permissive) or 25°C (non-permissive) for 3 days (or longer) prior to examination at the L4 stage. Other mutants were kept at 20°C except for *unc-26(s1710)* and *epg-9(bp320)* temperature experiments.

The penetrance of ATG-9 subsynaptic foci was quantified as percentage of animals showing subsynaptic foci of ATG-9:GFP at Zone 2. Mutant phenotype was defined as two or more than two subsynaptic foci of ATG-9::GFP at Zone 2 in endocytic mutants, or one or more than one subsynaptic foci in autophagy mutants. A Leica DM500B compound fluorescent microscope was used to visualize and screen the worm in different genetic backgrounds (Figures 2N, S2A, S2E, 4G-4H, 6J, 6P).

#### Expressivity of ATG-9::GFP Subsynaptic Foci at the Presynaptic Region

To quantify expressivity of ATG-9 subsynaptic foci at the presynaptic region (Zone 2), we obtained the plot profiles for individual presynaptic region (Zone 2) through segmented line scans using Fiji. An algorithm in MATLAB was designed to identify peaks along the line scans of Zone 2. The index of ATG-9 mislocalization at presynaptic region is defined as the number of peaks divided by the width of peaks at Zone 2 in each individual animal. In endocytic mutants, two populations of mutant phenotype were identified: ATG-9 is dim and diffuse at Zone 2; ATG-9 displays subsynaptic foci at Zone 2. Only endocytic mutants that display subsynaptic foci were quantified for the expressivity (Figure 2K).

Code available: https://github.com/yangsisi76/Quantify-distribution-of-cell-structures.

#### Mean Intensity of ATG-9 at Zone 2

To measure the level of ATG-9 at presynaptic regions, we obtained the fluorescent value for individual presynaptic region (Zone 2) through segmented line scans using Fiji. All settings for the confocal microscope and camera were kept identical between the wild type and *unc-26(s1710)* mutants. Mean fluorescent value of animals in the two genotypes was calculated by Fiji (Figure S2B).

#### RAB-3 and SNG-1 Enrichment at Zone 3

To quantify the distribution and enrichment of synaptic proteins, such as RAB-3 and SNG-1, we used methods as described (Dittman and Kaplan, 2006, Jang et al., 2016). Briefly, fluorescent values for the RAB-3 (SNG-1) in AIY neurons were obtained through segmented line scans using Fiji. A sliding window of 2μm was used to identify all the fluorescent peak and trough values for Zone 3 in each individual neuron. The synaptic enrichment was calculated as %*ΔF/F* as described (Jang et al., 2016, Dittman and Kaplan, 2006, Bai et al., 2010). In short, all the identified fluorescent peak and trough values (*F_peak_* and *F_trough_*) were averaged and used to calculate the %*ΔF/F* (100 x (*F_peak_* - *F_trough_*)/ *F_trough_*) (Bai et al., 2010, Dittman and Kaplan, 2006, Jang et al., 2016). All settings for the confocal microscope and camera were kept identical between the wild type and *unc-26(s1710)* mutants (Figures S3F and S5B).

#### Percentage of ATG-9 Subsynaptic Foci that Colocalize with the Subsynaptic Structures

To quantify the percentage of ATG-9 subsynaptic foci that colocalize with the subsynaptic structures (LGG-1 and CHC-1), we observed the transgenic lines *olaex1360;olaIs44* (pttx-3::gfp::lgg-1, pttx-3::mCherry::atg-9) and *olaex4290;olaIs34* (pttx-3::bfp::chc-1, pttx-3::atg-9::gfp). Confocal maximal projections were used, and percentage colocalization was calculated as the percentage of ATG-9 subsynaptic foci that colocalize with examined organelle markers (Figure 3M).

#### Activity-dependent Autophagy

To measure the synaptic autophagosomes in AIY, animals with *olaIs35* in the wild-type, *unc-13(e450)*, *atg-9(wy56)*, *unc-26(s1710)* and *unc-26(s1710);atg-9(wy56)* mutant backgrounds were grown in 20°C and then shifted to 25°C for 4 hours and assessed for number of LGG-1 puncta in the neurite of AIY under a Leica DM 5000B compound microscope (Hill et al., 2019). To show the comparison between 20°C and 25°C, the number of LGG-1 puncta at 25°C in each genotype was normalized by the number at 20°C (Figure 4L). Results before normalization are reported in Figure S4C.

#### Immuno-EM

To quantify the distribution of ATG-9 positive particles, animals with *olaex2264* (*punc-14::atg-9::GFP*) in the wild type were used. Quantifications were performed in FIJI. Cross-sectional area: an image of a 40nm section. Particles were counted using the Cell Counter plugin. Occasionally, a Gaussian Blur processing filter was applied in FIJI to help visualize structures in the image. Staining specificity, calculated as a signal to noise ratio of particle density in neuronal tissue divided by particle density in nearby *E coli* in the section, was >20 for all samples. Particles were considered localized near vesicles, plasma membrane, or mitochondria if the distance from the gold particle to that structure was <20 nm. This distance was chosen based on estimates of the size of GFP, a GFP antibody, and protein A crystal structures. To measure the area occupied by vesicles, a freehand shape was drawn around an apparent cluster of vesicles occupying most or all vesicles in the neurite. To measure area occupied by plasma membrane, the perimeter of the neurite was multiplied by 40, which accounts for the rough measured width of the plasma membrane plus the 20 nm radius in which a particle may be localized nearby (Figures 1G, S1A-S1D).

### Quantification of Immunoreactivity in Hippocampal Neuron Culture and GFP::CHC-1 in *C. elegans*

Quantification of ATG9 and CHC-1 clustering was performed using Fiji as previously described (Cao et al., 2020). Briefly, the same threshold was applied to all images after background subtraction and then ‘analyze particles’ function of Fiji was used to obtain the raw intensity values of masked regions (Figures S3G and 6I).

### Statistical Analysis

Statistical analysis and data plotting were conducted with Prism 7 software. We used Fisher’s exact test to determine statistical significance of categorical data in contingency table, such as the penetrance of ATG-9 phenotype in AIY. 95% confidence intervals were calculated by Wilson/Brown method and used for error bars. For the continuous data, ordinary one-way ANOVA with Tukey’s multiple comparisons test, Welch’s t test and Student’s t test were used to determine its statistical significance. The error bar represents the standard error of the mean (SEM). The p value for significant differences is reported in the figure legends.

## Reference

Alberti, A., Michelet, X., Djeddi, A. & Legouis, R. 2010. The autophagosomal protein LGG-2 acts synergistically with LGG-1 in dauer formation and longevity in C. elegans. Autophagy, 6, 622–33.

Alegre-Abarrategui, J. & Wade-Martins, R. 2009. Parkinson disease, LRRK2 and the endocytic-autophagic pathway. Autophagy, 5, 1208-10.

Anglade, P., Vyas, S., Javoy-Agid, F., Herrero, M. T., Michel, P. P., Marquez, J., Mouatt-Prigent, A., Ruberg, M., Hirsch, E. C. & Agid, Y. 1997. Apoptosis and autophagy in nigral neurons of patients with Parkinson’s disease. Histol Histopathol, 12, 25–31.

Azarnia Tehran, D., Kuijpers, M. & Haucke, V. 2018. Presynaptic endocytic factors in autophagy and neurodegeneration. Curr Opin Neurobiol, 48, 153–159.

Badolato, R. & Parolini, S. 2007. Novel insights from adaptor protein 3 complex deficiency. J Allergy Clin Immunol, 120, 735–41; quiz 742-3.

Bai, J. H., Hu, Z. T., Dittman, J. S., Pym, E. C. G. & Kaplan, J. M. 2010. Endophilin Functions as a Membrane-Bending Molecule and Is Delivered to Endocytic Zones by Exocytosis. Cell, 143, 430–441.

Bandres-Ciga, S., Saez-Atienzar, S., Bonet-Ponce, L., Billingsley, K., Vitale, D., Blauwendraat, C., Gibbs, J. R., Pihlstrom, L., Gan-Or, Z., International Parkinson’s Disease Genomics, C., Cookson, M. R., Nalls, M. A. & Singleton, A. B. 2019. The endocytic membrane trafficking pathway plays a major role in the risk of Parkinson’s disease. Mov Disord, 34, 460-468.

Binotti, B., Pavlos, N. J., Riedel, D., Wenzel, D., Vorbruggen, G., Schalk, A. M., Kuhnel, K., Boyken, J., Erck, C., Martens, H., Chua, J. J. & Jahn, R. 2015. The GTPase Rab26 links synaptic vesicles to the autophagy pathway. Elife, 4, e05597.

Biron, D., Shibuya, M., Gabel, C., Wasserman, S. M., Clark, D. A., Brown, A., Sengupta, P. & Samuel, A. D. 2006. A diacylglycerol kinase modulates long-term thermotactic behavioral plasticity in C. elegans. Nat Neurosci, 9, 1499–505.

Blauwendraat, C., Heilbron, K., Vallerga, C. L., Bandres-Ciga, S., Von Coelln, R., Pihlstrom, L., Simon-Sanchez, J., Schulte, C., Sharma, M., Krohn, L., Siitonen, A., Iwaki, H., Leonard, H., Noyce, A. J., Tan, M., Gibbs, J. R., Hernandez, D. G., Scholz, S. W., Jankovic, J., Shulman, L. M., Lesage, S., Corvol, J. C., Brice, A., Van Hilten, J. J., Marinus, J., Eerola-Rautio, J., Tienari, P., Majamaa, K., Toft, M., Grosset, D. G., Gasser, T., Heutink, P., Shulman, J. M., Wood, N., Hardy, J., Morris, H. R., Hinds, D. A., Gratten, J., Visscher, P. M., Gan-Or, Z., Nalls, M. A., Singleton, A. B., Team, A. R. & IPDGC 2019. Parkinson’s disease age at onset genome-wide association study: Defining heritability, genetic loci, and alpha-synuclein mechanisms. Movement Disorders, 34, 866–875.

Boyken, J., Gronborg, M., Riedel, D., Urlaub, H., Jahn, R. & Chua, J. J. E. 2013. Molecular Profiling of Synaptic Vesicle Docking Sites Reveals Novel Proteins but Few Differences between Glutamatergic and GABAergic Synapses. Neuron, 78, 285–297.

Brenner, S. 1974. The genetics of Caenorhabditis elegans. Genetics, 77, 71–94.

Bunge, M. B. 1973. Fine structure of nerve fibers and growth cones of isolated sympathetic neurons in culture. J Cell Biol, 56, 713-35.

Cao, M., Wu, Y., Ashrafi, G., Mccartney, A. J., Wheeler, H., Bushong, E. A., Boassa, D., Ellisman, M. H., Ryan, T. A. & De Camilli, P. 2017. Parkinson Sac Domain Mutation in Synaptojanin 1 Impairs Clathrin Uncoating at Synapses and Triggers Dystrophic Changes in Dopaminergic Axons. Neuron, 93, 882–896 e5.

Cao, M. A., Park, D., Wu, Y. M. & De Camilli, P. 2020. Absence of Sac2/INPP5F enhances the phenotype of a Parkinson’s disease mutation of synaptojanin 1. Proceedings of the National Academy of Sciences of the United States of America, 117, 12428–12434.

Chantranupong, L., Saulnier, J. L., Wang, W., Jones, D. R., Pacold, M. E. & Sabatini, B. L. 2020. Rapid purification and metabolomic profiling of synaptic vesicles from mammalian brain. Elife, 9.

Cheung, Z. H. & Ip, N. Y. 2009. The emerging role of autophagy in Parkinson’s disease. Mol Brain, 2, 29.

Clark, D. A., Biron, D., Sengupta, P. & Samuel, A. D. T. 2006. The AFD sensory neurons encode multiple functions underlying thermotactic behavior in Caenorhabditis elegans. Journal of Neuroscience, 26, 7444–7451.

Colon-Ramos, D. A., Margeta, M. A. & Shen, K. 2007. Glia promote local synaptogenesis through UNC-6 (netrin) signaling in C-elegans. Science, 318, 103–106.

Crawley, O., Opperman, K. J., Desbois, M., Adrados, I., Borgen, M. A., Giles, A. C., Duckett, D. R. & Grill, B. 2019. Autophagy is inhibited by ubiquitin ligase activity in the nervous system. Nature Communications, 10.

Cremona, O., Di Paolo, G., Wenk, M. R., Luthi, A., Kim, W. T., Takei, K., Daniell, L., Nemoto, Y., Shears, S. B., Flavell, R. A., Mccormick, D. A. & De Camilli, P. 1999. Essential role of phosphoinositide metabolism in synaptic vesicle recycling. Cell, 99, 179–88.

De Pace, R., Skirzewski, M., Damme, M., Mattera, R., Mercurio, J., Foster, A. M., Cuitino, L., Jarnik, M., Hoffmann, V., Morris, H. D., Han, T. U., Mancini, G. M. S., Buonanno, A. & Bonifacino, J. S. 2018. Altered distribution of ATG9A and accumulation of axonal aggregates in neurons from a mouse model of AP-4 deficiency syndrome. PLoS Genet, 14, e1007363.

Dell’Angelica, E. C. 2009. AP-3-dependent trafficking and disease: the first decade. Curr Opin Cell Biol, 21, 552–9.

Dell’Angelica, E. C., Ohno, H., Ooi, C. E., Rabinovich, E., Roche, K. W. & Bonifacino, J. S. 1997. AP-3: an adaptor-like protein complex with ubiquitous expression. EMBO J, 16, 917–28.

Dittman, J. S. & Kaplan, J. M. 2006. Factors regulating the abundance and localization of synaptobrevin in the plasma membrane. Proc Natl Acad Sci U S A, 103, 11399–404.

Feng, Y. C. & Klionsky, D. J. 2017. Autophagic membrane delivery through ATG9. Cell Research, 27, 161–162.

Ferguson, S. M., Brasnjo, G., Hayashi, M., Wolfel, M., Collesi, C., Giovedi, S., Raimondi, A., Gong, L. W., Ariel, P., Paradise, S., O’Toole, E., Flavell, R., Cremona, O., Miesenbock, G., Ryan, T. A. & De Camilli, P. 2007. A selective activity-dependent requirement for dynamin 1 in synaptic vesicle endocytosis. Science, 316, 570–574.

Ferguson, S. M. & De Camilli, P. 2012. Dynamin, a membrane-remodelling GTPase. Nat Rev Mol Cell Biol, 13, 75–88.

Ferguson, S. M., Raimondi, A., Paradise, S., Shen, H., Mesaki, K., Ferguson, A., Destaing, O., Ko, G., Takasaki, J., Cremona, O., E, O. T. & De Camilli, P. 2009. Coordinated actions of actin and BAR proteins upstream of dynamin at endocytic clathrin-coated pits. Dev Cell, 17, 811–22.

Gan, Q. & Watanabe, S. 2018. Synaptic Vesicle Endocytosis in Different Model Systems. Front Cell Neurosci, 12, 171.

George, A. A., Hayden, S., Stanton, G. R. & Brockerhoff, S. E. 2016. Arf6 and the 5’phosphatase of synaptojanin 1 regulate autophagy in cone photoreceptors. Bioessays, 38 Suppl 1, S119–35.

Gomez-Sanchez, R., Rose, J., Guimaraes, R., Mari, M., Papinski, D., Rieter, E., Geerts, W. J., Hardenberg, R., Kraft, C., Ungermann, C. & Reggiori, F. 2018. Atg9 establishes Atg2-dependent contact sites between the endoplasmic reticulum and phagophores. J Cell Biol, 217, 2743–2763.

Guardia, C. M., Christenson, E. T., Zhou, W., Tan, X. F., Lian, T., Faraldo-Gomez, J. D., Bonifacino, J. S., Jiang, J. & Banerjee, A. 2020. The structure of human ATG9A and its interplay with the lipid bilayer. Autophagy, 1-2.

Harris, T. W., Hartwieg, E., Horvitz, H. R. & Jorgensen, E. M. 2000. Mutations in synaptojanin disrupt synaptic vesicle recycling. Journal of Cell Biology, 150, 589–599.

Hata, Y., Slaughter, C. A. & Sudhof, T. C. 1993. Synaptic vesicle fusion complex contains unc-18 homologue bound to syntaxin. Nature, 366, 347–51.

Hawk, J. D., Calvo, A. C., Liu, P., Almoril-Porras, A., Aljobeh, A., Torruella-Suarez, M. L., Ren, I., Cook, N., Greenwood, J., Luo, L. J., Wang, Z. W., Samuel, A. D. T. & Colon-Ramos, D. A. 2018. Integration of Plasticity Mechanisms within a Single Sensory Neuron of C. elegans Actuates a Memory. Neuron, 97, 356-+.

Hayashi, M., Raimondi, A., O’Toole, E., Paradise, S., Collesi, C., Cremona, O., Ferguson, S. M. & De Camilli, P. 2008. Cell- and stimulus-dependent heterogeneity of synaptic vesicle endocytic recycling mechanisms revealed by studies of dynamin 1-null neurons. Proc Natl Acad Sci U S A, 105, 2175–80.

Hill, S. E., Kauffman, K. J., Krout, M., Richmond, J. E., Melia, T. J. & Colon-Ramos, D. A. 2019. Maturation and Clearance of Autophagosomes in Neurons Depends on a Specific Cysteine Protease Isoform, ATG-4.2. Developmental Cell, 49, 251-+.

Hoffmann, S., Orlando, M., Andrzejak, E., Bruns, C., Trimbuch, T., Rosenmund, C., Garner, C. C. & Ackermann, F. 2019. Light-Activated ROS Production Induces Synaptic Autophagy. J Neurosci, 39, 2163–2183.

Huang, S., Jia, K., Wang, Y., Zhou, Z. & Levine, B. 2013. Autophagy genes function in apoptotic cell corpse clearance during C. elegans embryonic development. Autophagy, 9, 138–149.

Jang, S., Nelson, J. C., Bend, E. G., Rodriguez-Laureano, L., Tueros, F. G., Cartagenova, L., Underwood, K., Jorgensen, E. M. & Colon-Ramos, D. A. 2016. Glycolytic Enzymes Localize to Synapses under Energy Stress to Support Synaptic Function. Neuron, 90, 278–291.

Jang, S., Xuan, Z., Lagoy, R. C., Jawerth, L. M., Gonzalez, I. J., Singh, M., Prashad, S., Kim, H. S., Patel, A., Albrecht, D. R., Hyman, A. A. & Colon-Ramos, D. A. 2020. Phosphofructokinase Relocalizes into Subcellular Compartments with Liquid-like Properties In Vivo. Biophys J. Karabiyik, C., Lee, M. J. & Rubinsztein, D. C. 2017. Autophagy impairment in Parkinson’s disease. Essays Biochem, 61, 711-720.

Karanasios, E., Walker, S. A., Okkenhaug, H., Manifava, M., Hummel, E., Zimmermann, H., Ahmed, Q., Domart, M. C., Collinson, L. & Ktistakis, N. T. 2016. Autophagy initiation by ULK complex assembly on ER tubulovesicular regions marked by ATG9 vesicles. Nature Communications, 7.

Katsumata, K., Nishiyama, J., Inoue, T., Mizushima, N., Takeda, J. & Yuzaki, M. 2010. Dynein- and activity-dependent retrograde transport of autophagosomes in neuronal axons. Autophagy, 6, 378–85.

Kim, W. T., Chang, S. H., Daniell, L., Cremona, O., Di Paolo, G. & De Camilli, P. 2002. Delayed reentry of recycling vesicles into the fusion-competent synaptic vesicle pool in synaptojanin 1 knockout mice. Proceedings of the National Academy of Sciences of the United States of America, 99, 17143–17148.

Kolotuev, I. 2014. Positional correlative anatomy of invertebrate model organisms increases efficiency of TEM data production. Microsc Microanal, 20, 1392–403.

Kolotuev, I., Bumbarger, D. J., Labouesse, M. & Schwab, Y. 2012. Targeted ultramicrotomy: a valuable tool for correlated light and electron microscopy of small model organisms. Methods Cell Biol, 111, 203–22.

Kononenko, N. L., Classen, G. A., Kuijpers, M., Puchkov, D., Maritzen, T., Tempes, A., Malik, A. R., Skalecka, A., Bera, S., Jaworski, J. & Haucke, V. 2017. Retrograde transport of TrkB-containing autophagosomes via the adaptor AP-2 mediates neuronal complexity and prevents neurodegeneration. Nat Commun, 8, 14819.

Krebs, C. E., Karkheiran, S., Powell, J. C., Cao, M., Makarov, V., Darvish, H., Di Paolo, G., Walker, R. H., Shahidi, G. A., Buxbaum, J. D., De Camilli, P., Yue, Z. Y. & Paisan-Ruiz, C. 2013. The Sac1 Domain of SYNJ1 Identified Mutated in a Family with Early-Onset Progressive Parkinsonism with Generalized Seizures. Human Mutation, 34, 1200–1207.

Kulkarni, A., Chen, J. & Maday, S. 2018. Neuronal autophagy and intercellular regulation of homeostasis in the brain. Curr Opin Neurobiol, 51, 29–36.

Liang, Q. Q., Yang, P. G., Tian, E., Han, J. H. & Zhang, H. 2012. The C. elegans ATG101 homolog EPG-9 directly interacts with EPG-1/Atg13 and is essential for autophagy. Autophagy, 8, 1426–1433.

Liang, Y. & Sigrist, S. 2018. Autophagy and proteostasis in the control of synapse aging and disease. Curr Opin Neurobiol, 48, 113–121.

Lu, Q., Yang, P., Huang, X., Hu, W., Guo, B., Wu, F., Lin, L., Kovacs, A. L., Yu, L. & Zhang, H. 2011. The WD40 repeat PtdIns(3)P-binding protein EPG-6 regulates progression of omegasomes to autophagosomes. Dev Cell, 21, 343–57.

Lynch-Day, M. A., Mao, K., Wang, K., Zhao, M. & Klionsky, D. J. 2012. The role of autophagy in Parkinson’s disease. Cold Spring Harb Perspect Med, 2, a009357.

Maday, S., Wallace, K. E. & Holzbaur, E. L. 2012. Autophagosomes initiate distally and mature during transport toward the cell soma in primary neurons. J Cell Biol, 196, 407–17.

Maeda, S., Yamamoto, H., Kinch, L. N., Garza, C. M., Takahashi, S., Otomo, C., Grishin, N. V., Forli, S., Mizushima, N. & Otomo, T. 2020. Structure, lipid scrambling activity and role in autophagosome formation of ATG9A. Nat Struct Mol Biol.

Manil-Segalen, M., Lefebvre, C., Jenzer, C., Trichet, M., Boulogne, C., Satiat-Jeunemaitre, B. & Legouis, R. 2014. The C. elegans LC3 acts downstream of GABARAP to degrade autophagosomes by interacting with the HOPS subunit VPS39. Dev Cell, 28, 43–55.

Manning, L. & Richmond, J. 2015. High-Pressure Freeze and Freeze Substitution Electron Microscopy in C. elegans. Methods Mol Biol, 1327, 121–40.

Matoba, K., Kotani, T., Tsutsumi, A., Tsuji, T., Mori, T., Noshiro, D., Sugita, Y., Nomura, N., Iwata, S., Ohsumi, Y., Fujimoto, T., Nakatogawa, H., Kikkawa, M. & Noda, N. N. 2020. Atg9 is a lipid scramblase that mediates autophagosomal membrane expansion. Nat Struct Mol Biol.

Matoba, K. & Noda, N. N. 2020. Secret of Atg9: lipid scramblase activity drives de novo autophagosome biogenesis. Cell Death Differ.

Mattera, R., Park, S. Y., De Pace, R., Guardia, C. M. & Bonifacino, J. S. 2017. AP-4 mediates export of ATG9A from the trans-Golgi network to promote autophagosome formation. Proc Natl Acad Sci U S A, 114, E10697–E10706.

Menzies, F. M., Fleming, A., Caricasole, A., Bento, C. F., Andrews, S. P., Ashkenazi, A., Fullgrabe, J., Jackson, A., Jimenez Sanchez, M., Karabiyik, C., Licitra, F., Lopez Ramirez, A., Pavel, M., Puri, C., Renna, M., Ricketts, T., Schlotawa, L., Vicinanza, M., Won, H., Zhu, Y., Skidmore, J. & Rubinsztein, D. C. 2017. Autophagy and Neurodegeneration: Pathogenic Mechanisms and Therapeutic Opportunities. Neuron, 93, 1015–1034.

Menzies, F. M., Fleming, A. & Rubinsztein, D. C. 2015. Compromised autophagy and neurodegenerative diseases. Nat Rev Neurosci, 16, 345–57.

Milosevic, I., Giovedi, S., Lou, X., Raimondi, A., Collesi, C., Shen, H., Paradise, S., O’Toole, E., Ferguson, S., Cremona, O. & De Camilli, P. 2011. Recruitment of endophilin to clathrin-coated pit necks is required for efficient vesicle uncoating after fission. Neuron, 72, 587–601.

Murdoch, J. D., Rostosky, C. M., Gowrisankaran, S., Arora, A. S., Soukup, S. F., Vidal, R., Capece, V., Freytag, S., Fischer, A., Verstreken, P., Bonn, S., Raimundo, N. & Milosevic, I. 2016. Endophilin-A Deficiency Induces the Foxo3a-Fbxo32 Network in the Brain and Causes Dysregulation of Autophagy and the Ubiquitin-Proteasome System. Cell Rep, 17, 1071–1086.

Nakatsu, F. & Ohno, H. 2003. Adaptor protein complexes as the key regulators of protein sorting in the post-Golgi network. Cell Structure and Function, 28, 419–429.

Nalls, M. A., Pankratz, N., Lill, C. M., Do, C. B., Hernandez, D. G., Saad, M., Destefano, A. L., Kara, E., Bras, J., Sharma, M., Schulte, C., Keller, M. F., Arepalli, S., Letson, C., Edsall, C., Stefansson, H., Liu, X., Pliner, H., Lee, J. H., Cheng, R., International Parkinson’s Disease Genomics, C., Parkinson’s Study Group Parkinson’s Research: The Organized, G. I., Andme, Genepd, Neurogenetics Research, C., Hussman Institute Of Human, G., Ashkenazi Jewish Dataset, I., Cohorts For, H., Aging Research In Genetic, E., North American Brain Expression, C., United Kingdom Brain Expression, C., Greek Parkinson’s Disease, C., Alzheimer Genetic Analysis, G., Ikram, M. A., Ioannidis, J. P., Hadjigeorgiou, G. M., Bis, J. C., Martinez, M., Perlmutter, J. S., Goate, A., Marder, K., Fiske, B., Sutherland, M., Xiromerisiou, G., Myers, R. H., Clark, L. N., Stefansson, K., Hardy, J. A., Heutink, P., Chen, H., Wood, N. W., Houlden, H., Payami, H., Brice, A., Scott, W. K., Gasser, T., Bertram, L., Eriksson, N., Foroud, T. & Singleton, A. B. 2014. Large-scale meta-analysis of genome-wide association data identifies six new risk loci for Parkinson’s disease. Nat Genet, 46, 989-93.

Noda, T. 2017. Autophagy in the context of the cellular membrane-trafficking system: the enigma of Atg9 vesicles. Biochemical Society Transactions, 45, 1323-1331.

Ohashi, Y. & Munro, S. 2010. Membrane Delivery to the Yeast Autophagosome from the Golgi-Endosomal System. Molecular Biology of the Cell, 21, 3998–4008.

Paix, A., Folkmann, A. & Seydoux, G. 2017. Precision genome editing using CRISPR-Cas9 and linear repair templates in C. elegans. Methods, 121, 86–93.

Park, S. Y. & Guo, X. 2014. Adaptor protein complexes and intracellular transport. Biosci Rep, 34.

Popovic, D. & Dikic, I. 2014. TBC1D5 and the AP2 complex regulate ATG9 trafficking and initiation of autophagy. EMBO Rep, 15, 392–401.

Puri, C., Renna, M., Bento, C. F., Moreau, K. & Rubinsztein, D. C. 2013. Diverse autophagosome membrane sources coalesce in recycling endosomes. Cell, 154, 1285–99.

Puri, C., Renna, M., Bento, C. F., Moreau, K. & Rubinsztein, D. C. 2014. ATG16L1 meets ATG9 in recycling endosomes: additional roles for the plasma membrane and endocytosis in autophagosome biogenesis. Autophagy, 10, 182–4.

Quadri, M., Fang, M., Picillo, M., Olgiati, S., Breedveld, G. J., Graafland, J., Wu, B., Xu, F., Erro, R., Amboni, M., Pappata, S., Quarantelli, M., Annesi, G., Quattrone, A., Chien, H. F., Barbosa, E. R., International Parkinsonism Genetics, N., Oostra, B. A., Barone, P., Wang, J. & Bonifati, V. 2013. Mutation in the SYNJ1 gene associated with autosomal recessive, early-onset Parkinsonism. Hum Mutat, 34, 1208–15.

Raimondi, A., Ferguson, S. M., Lou, X., Armbruster, M., Paradise, S., Giovedi, S., Messa, M., Kono, N., Takasaki, J., Cappello, V., O’Toole, E., Ryan, T. A. & De Camilli, P. 2011. Overlapping role of dynamin isoforms in synaptic vesicle endocytosis. Neuron, 70, 1100–14.

Reggiori, F., Shintani, T., Nair, U. & Klionsky, D. J. 2005. Atg9 cycles between mitochondria and the pre-autophagosomal structure in yeasts. Autophagy, 1, 101–109.

Reggiori, F., Tucker, K. A., Stromhaug, P. E. & Klionsky, D. J. 2004. The Atg1-Atg13 complex regulates Atg9 and Atg23 retrieval transport from the pre-autophagosomal structure. Developmental Cell, 6, 79–90.

Richmond, J. E., Davis, W. S. & Jorgensen, E. M. 1999. UNC-13 is required for synaptic vesicle fusion in C-elegans. Nature Neuroscience, 2, 959–964.

Rostaing, P., Weimer, R. M., Jorgensen, E. M., Triller, A. & Bessereau, J. L. 2004. Preservation of immunoreactivity and fine structure of adult C. elegans tissues using high-pressure freezing. J Histochem Cytochem, 52, 1–12.

Rout, M. P. & Field, M. C. 2017. The Evolution of Organellar Coat Complexes and Organization of the Eukaryotic Cell. Annu Rev Biochem, 86, 637–657.

Saheki, Y. & De Camilli, P. 2012. Synaptic vesicle endocytosis. Cold Spring Harb Perspect Biol, 4, a005645.

Sawa-Makarska, J., Baumann, V., Coudevylle, N., Von Bulow, S., Nogellova, V., Abert, C., Schuschnig, M., Graef, M., Hummer, G. & Martens, S. 2020. Reconstitution of autophagosome nucleation defines Atg9 vesicles as seeds for membrane formation. Science, 369.

Schindelin, J., Arganda-Carreras, I., Frise, E., Kaynig, V., Longair, M., Pietzsch, T., Preibisch, S., Rueden, C., Saalfeld, S., Schmid, B., Tinevez, J. Y., White, D. J., Hartenstein, V., Eliceiri, K., Tomancak, P. & Cardona, A. 2012. Fiji: an open-source platform for biological-image analysis. Nat Methods, 9, 676–82.

Schreij, A. M., Fon, E. A. & Mcpherson, P. S. 2016. Endocytic membrane trafficking and neurodegenerative disease. Cell Mol Life Sci, 73, 1529–45.

Sekito, T., Kawamata, T., Ichikawa, R., Suzuki, K. & Ohsumi, Y. 2009. Atg17 recruits Atg9 to organize the pre-autophagosomal structure. Genes Cells, 14, 525–38.

Shehata, M., Matsumura, H., Okubo-Suzuki, R., Ohkawa, N. & Inokuchi, K. 2012. Neuronal Stimulation Induces Autophagy in Hippocampal Neurons That Is Involved in AMPA Receptor Degradation after Chemical Long-Term Depression. Journal of Neuroscience, 32, 10413–10422.

Son, J. H., Shim, J. H., Kim, K. H., Ha, J. Y. & Han, J. Y. 2012. Neuronal autophagy and neurodegenerative diseases. Exp Mol Med, 44, 89–98.

Soukup, S. F., Kuenen, S., Vanhauwaert, R., Manetsberger, J., Hernandez-Diaz, S., Swerts, J., Schoovaerts, N., Vilain, S., Gounko, N. V., Vints, K., Geens, A., De Strooper, B. & Verstreken, P. 2016. A LRRK2-Dependent EndophilinA Phosphoswitch Is Critical for Macroautophagy at Presynaptic Terminals. Neuron, 92, 829–844.

Stavoe, A. K. H., Hill, S. E., Hall, D. H. & Colon-Ramos, D. A. 2016. KIF1A/UNC-104 Transports ATG-9 to Regulate Neurodevelopment and Autophagy at Synapses. Developmental Cell, 38, 171–185.

Stavoe, A. K. H. & Holzbaur, E. L. F. 2019. Axonal autophagy: Mini-review for autophagy in the CNS. Neurosci Lett, 697, 17–23.

Sudhof, T. C. 1995. The synaptic vesicle cycle: a cascade of protein-protein interactions. Nature, 375, 645–53.

Suzuki, K., Kirisako, T., Kamada, Y., Mizushima, N., Noda, T. & Ohsumi, Y. 2001. The pre-autophagosomal structure organized by concerted functions of APG genes is essential for autophagosome formation. EMBO J, 20, 5971–81.

Trinh, J. & Farrer, M. 2013. Advances in the genetics of Parkinson disease. Nature Reviews Neurology, 9, 445–454.

Tsukada, M. & Ohsumi, Y. 1993. Isolation and characterization of autophagy-defective mutants of Saccharomyces cerevisiae. FEBS Lett, 333, 169–74.

Van Der Vaart, A. & Reggiori, F. 2010. The Golgi complex as a source for yeast autophagosomal membranes. Autophagy, 6, 800–801.

Vanhauwaert, R., Kuenen, S., Masius, R., Bademosi, A., Manetsberger, J., Schoovaerts, N., Bounti, L., Gontcharenko, S., Swerts, J., Vilain, S., Picillo, M., Barone, P., Munshi, S. T., De Vrij, F. M., Kushner, S. A., Gounko, N. V., Mandemakers, W., Bonifati, V., Meunier, F. A., Soukup, S. F. & Verstreken, P. 2017. The SAC1 domain in synaptojanin is required for autophagosome maturation at presynaptic terminals. EMBO J, 36, 1392–1411.

Verstreken, P., Koh, T. W., Schulze, K. L., Zhai, R. G., Hiesinger, P. R., Zhou, Y., Mehta, S. Q., Cao, Y., Roos, J. & Bellen, H. J. 2003. Synaptojanin is recruited by Endophilin to promote synaptic vesicle uncoating. Neuron, 40, 733–748.

Vidyadhara, D. J., Lee, J. E. & Chandra, S. S. 2019. Role of the endolysosomal system in Parkinson’s disease. Journal of Neurochemistry, 150, 487–506.

Vijayan, V. & Verstreken, P. 2017. Autophagy in the presynaptic compartment in health and disease. J Cell Biol, 216, 1895–1906.

Wang, T., Martin, S., Papadopulos, A., Harper, C., Mavlyutov, T., Niranjan, D., Glass, N., Cooper-White, J., Sibarita, J. B., Choquet, D., Davletov, B. & Meunier, F. 2015. Control of autophagosome axonal retrograde flux by presynaptic activity unveiled using botulinum neurotoxin type-A. Journal of Neurochemistry, 134, 165–165.

Watanabe, S., Trimbuch, T., Camacho-Perez, M., Rost, B. R., Brokowski, B., Sohl-Kielczynski, B., Felies, A., Davis, M. W., Rosenmund, C. & Jorgensen, E. M. 2014. Clathrin regenerates synaptic vesicles from endosomes. Nature, 515, 228–33.

Webber, J. L., Young, A. R. J. & Tooze, S. A. 2007. Atg9 trafficking in mammalian cells. Autophagy, 3, 54–56.

White, J. G., Southgate, E., Thomson, J. N. & Brenner, S. 1986. The structure of the nervous system of the nematode Caenorhabditis elegans. Philos Trans R Soc Lond B Biol Sci, 314, 1–340.

Wu, F., Watanabe, Y., Guo, X. Y., Qi, X., Wang, P., Zhao, H. Y., Wang, Z., Fujioka, Y., Zhang, H., Ren, J. Q., Fang, T. C., Shen, Y. X., Feng, W., Hu, J. J., Noda, N. N. & Zhang, H. 2015. Structural Basis of the Differential Function of the Two C. elegans Atg8 Homologs, LGG-1 and LGG-2, in Autophagy. Mol Cell, 60, 914-29.

Yamamoto, H., Kakuta, S., Watanabe, T. M., Kitamura, A., Sekito, T., Kondo-Kakuta, C., Ichikawa, R., Kinjo, M. & Ohsumi, Y. 2012. Atg9 vesicles are an important membrane source during early steps of autophagosome formation. Journal of Cell Biology, 198, 219–233.

Yorimitsu, T. & Klionsky, D. J. 2005. Autophagy: molecular machinery for self-eating. Cell Death Differ, 12 Suppl 2, 1542–52.

Zavodszky, E., Seaman, M. N., Moreau, K., Jimenez-Sanchez, M., Breusegem, S. Y., Harbour, M. E. & Rubinsztein, D. C. 2014. Mutation in VPS35 associated with Parkinson’s disease impairs WASH complex association and inhibits autophagy. Nat Commun, 5, 3828.

Zhou, C., Ma, K., Gao, R., Mu, C., Chen, L., Liu, Q., Luo, Q., Feng, D., Zhu, Y. & Chen, Q. 2017. Regulation of mATG9 trafficking by Src- and ULK1-mediated phosphorylation in basal and starvation-induced autophagy. Cell Res, 27, 184–201.

