## supplemental figures for "Presynaptic autophagy is coupled to the synaptic vesicle cycle via ATG-9"

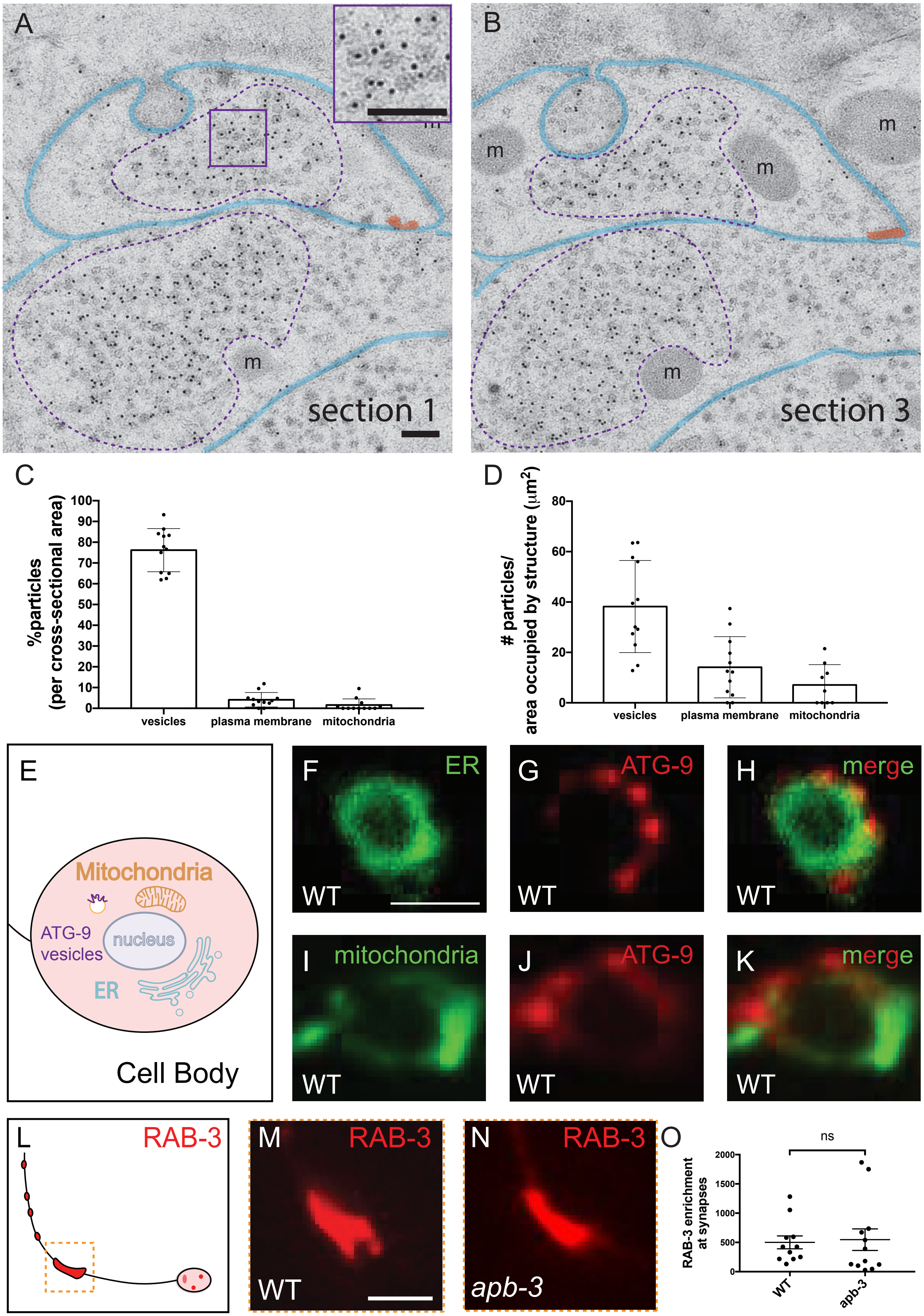


### Supplemental figure 1

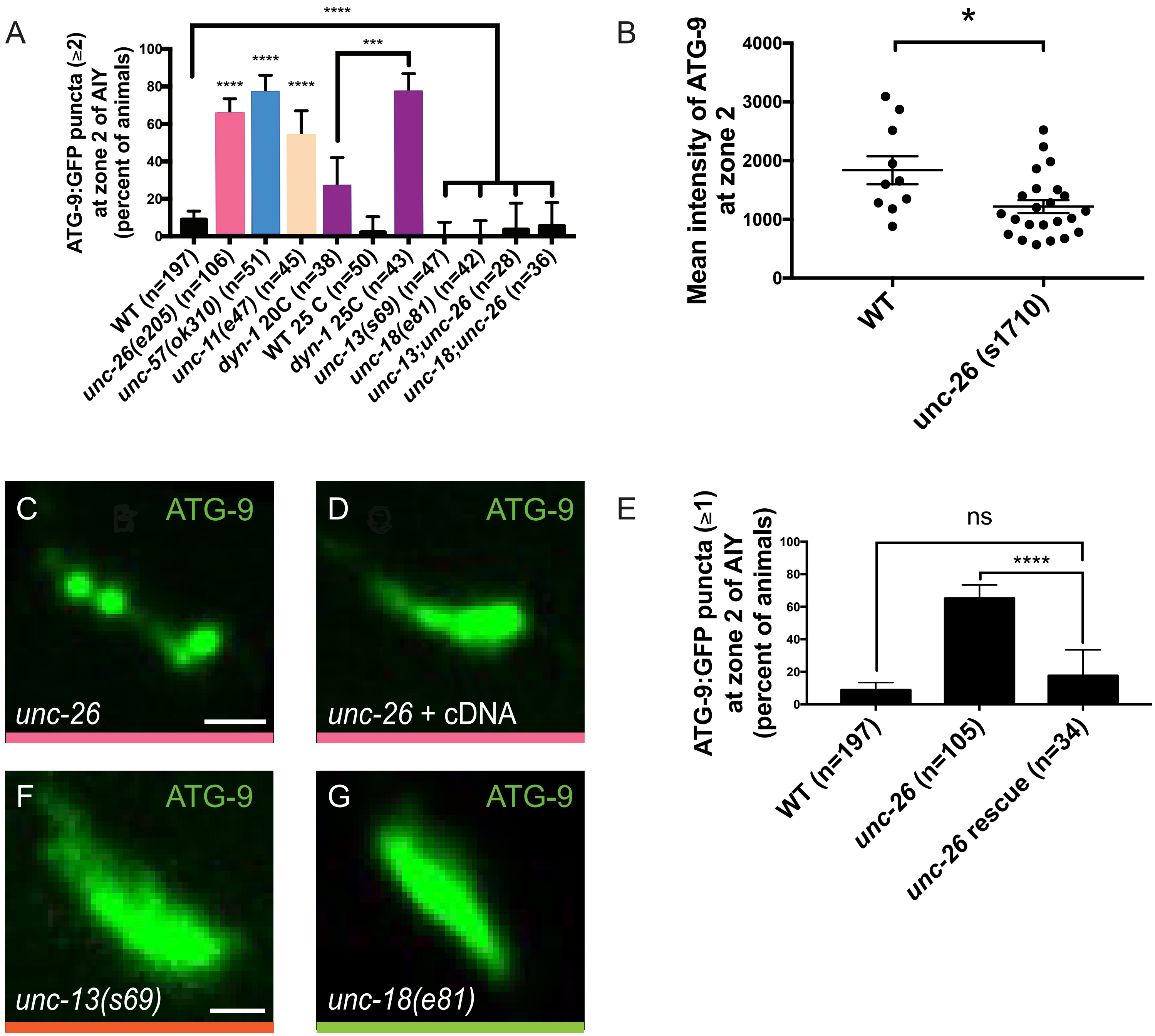


Supplemental figure 2

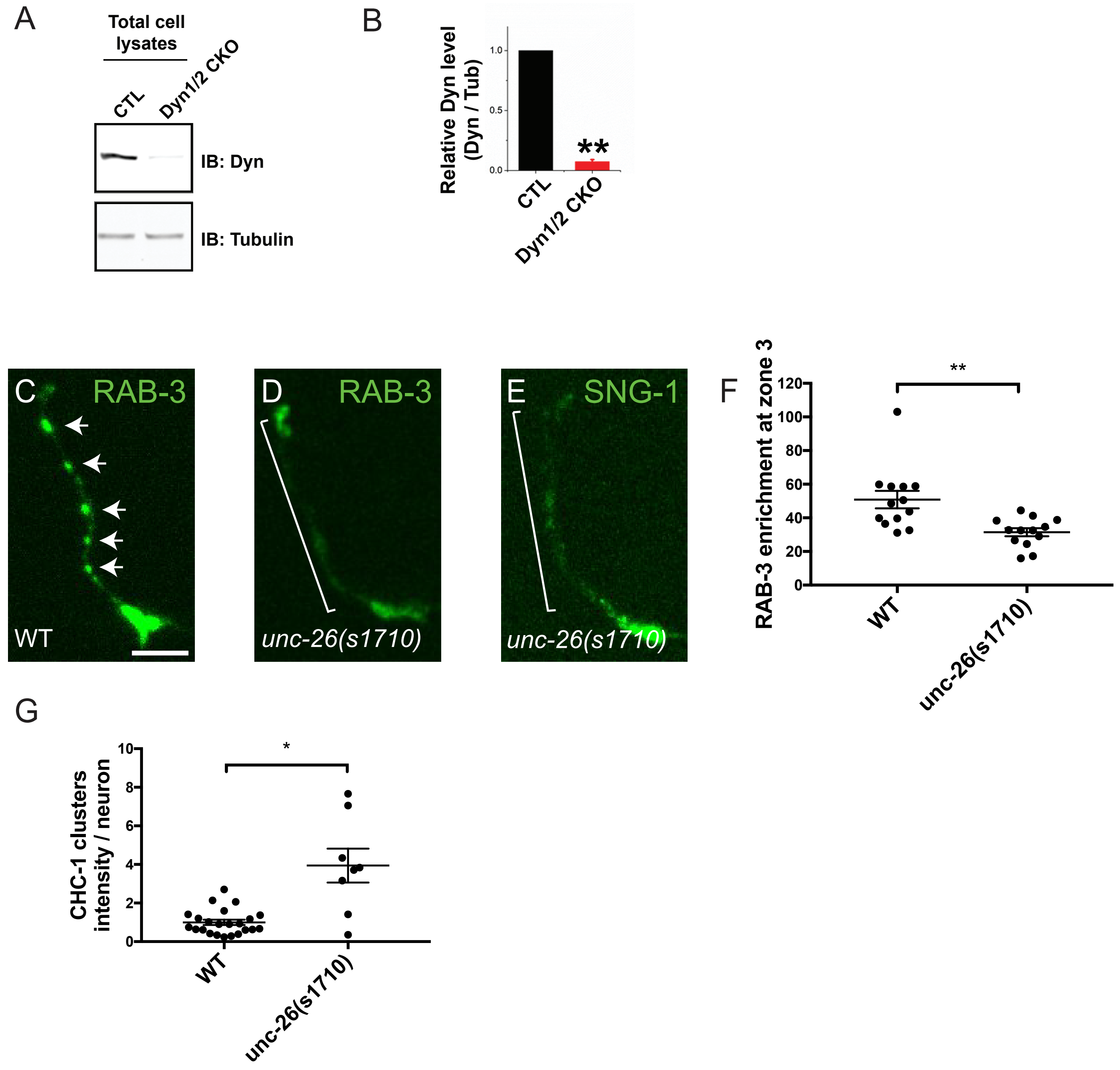


Supplemental figure 3

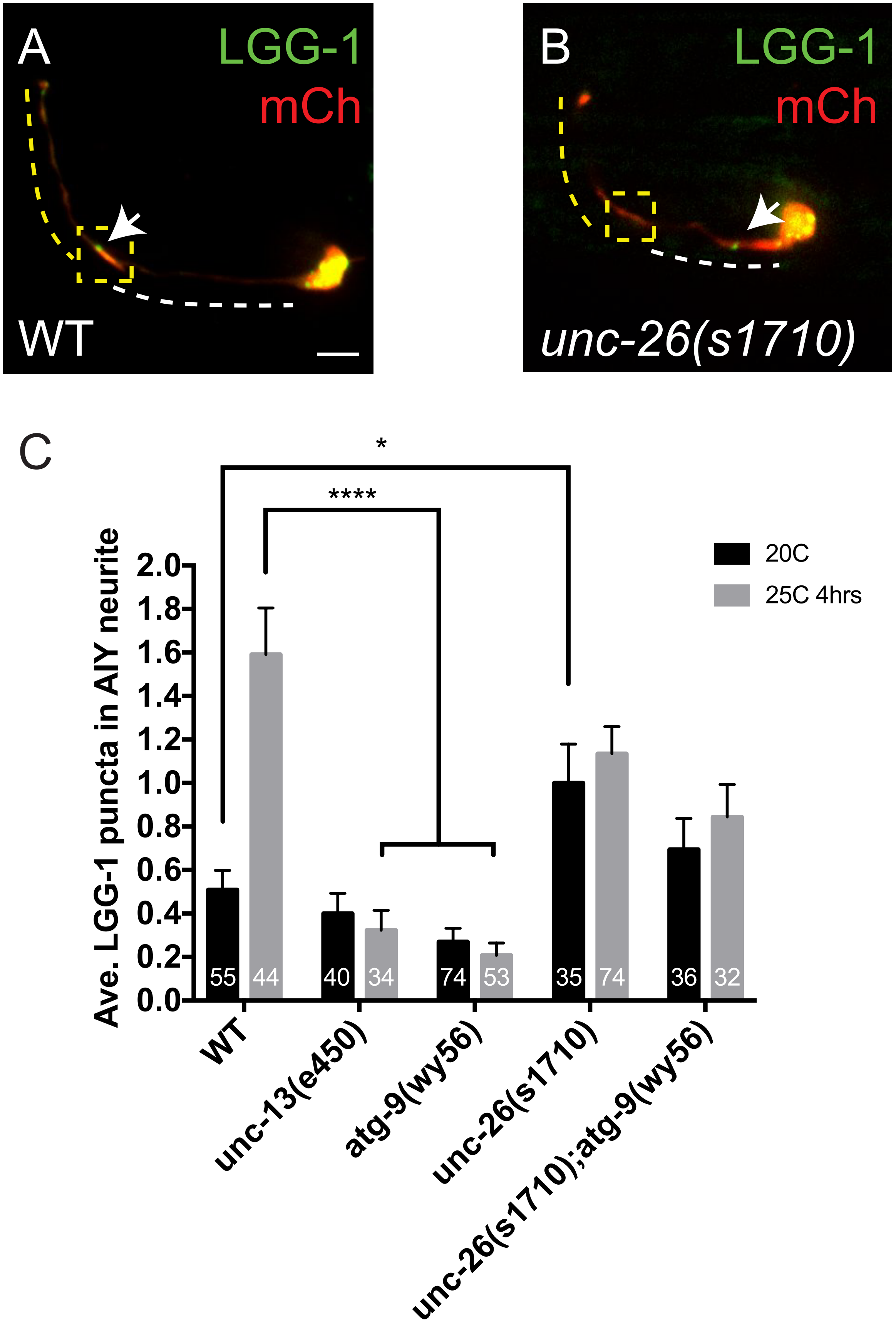


Supplemental figure 4

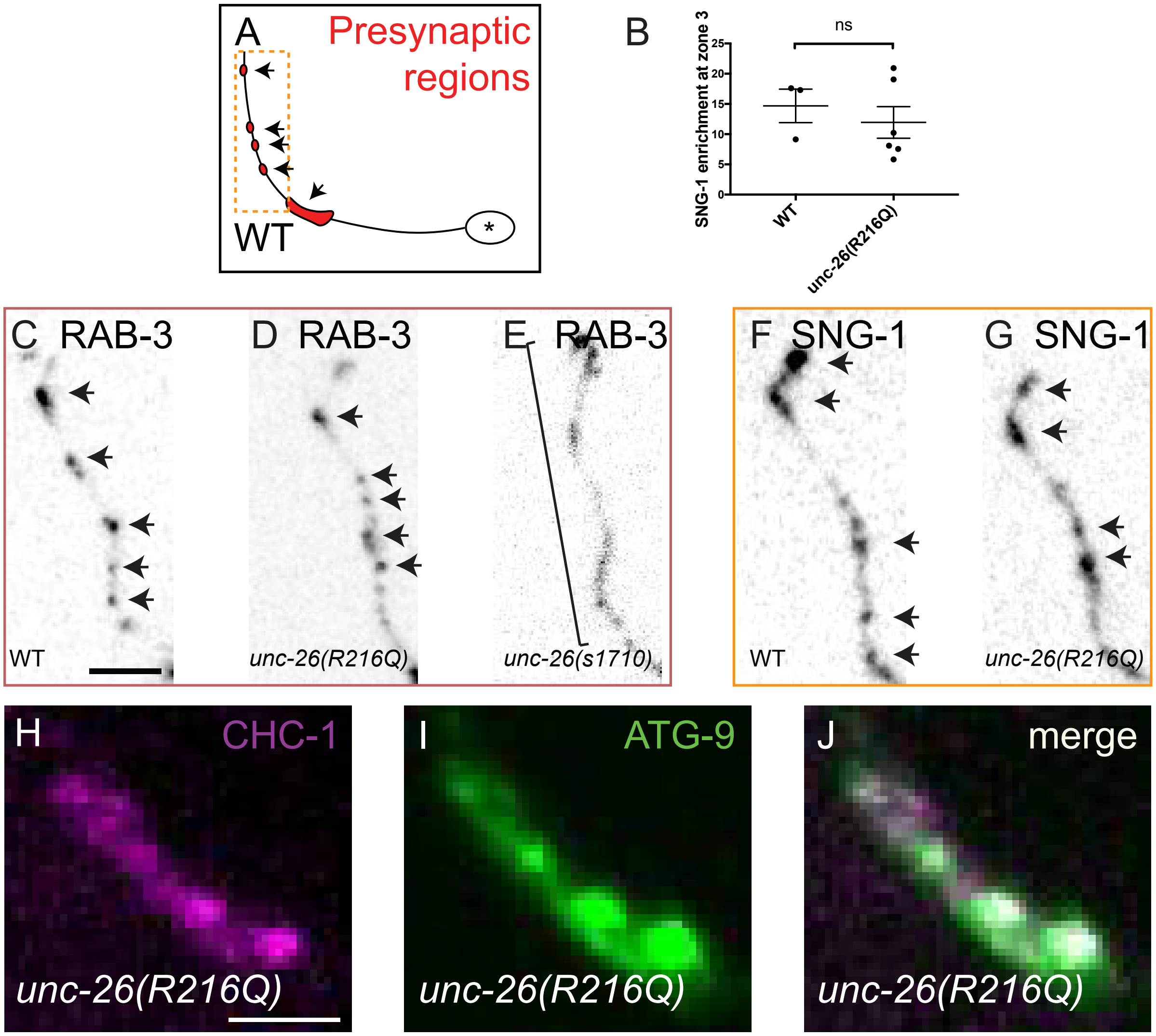


### Supplemental figure 5
